# Mapping the ultrastructural topology of the corynebacterial cell surface

**DOI:** 10.1101/2024.09.05.611374

**Authors:** Buse Isbilir, Anna Yeates, Vikram Alva, Tanmay A.M. Bharat

## Abstract

*Corynebacterium glutamicum* is a diderm bacterium extensively used in the industrial-scale production of amino acids. Corynebacteria belong to the bacterial family *Mycobacteriaceae*, which is characterized by a highly unusual cell envelope with an outer membrane consisting of mycolic acids. Despite the occurrence of this distinctive cell envelope in several bacterial pathogens, including *Corynebacterium diphtheriae, Mycobacterium tuberculosis*, and *Mycobacterium leprae*, its ultrastructural and molecular details remain elusive.

To address this, we investigated the cell envelope of *C. glutamicum* using electron cryotomography and cryomicroscopy of focused ion beam-milled cells. Our high-resolution images allowed us to accurately map the different components of the cell envelope into the tomographic density. Our data reveal that *C. glutamicum* has a variable cell envelope, with the outermost layer comprising the surface (S-)layer, which decorates the mycomembrane in a patchy manner. We further isolated and resolved the structure of the S-layer at 3.1 Å resolution using single particle electron cryomicroscopy. Our structure shows that the S-layer of *C. glutamicum* is composed of a hexagonal array of the PS2 protein, which interacts directly with the mycomembrane via a coiled coil-containing anchoring segment. Bioinformatic analyses revealed that the PS2 S-layer is sparsely yet exclusively present within the *Corynebacterium* genus and absent in other genera of the *Mycobacteriaceae* family, suggesting distinct evolutionary pathways in the development of their cell envelopes.

Our structural and cellular data collectively provide a high-resolution topography of the unusual *C. glutamicum* cell surface, features of which are shared by many pathogenic and microbiome-associated bacteria, as well as by several industrially significant bacterial species. This study, therefore, provides a strong experimental framework for understanding cell envelopes that contain mycolic acids.

## Introduction

Cell envelopes serve as primary interfaces between cells and their surroundings. Cell envelopes protect the internal components of the cell (Albers & Meyer, 2011; Bharat et al., 2021), regulate cell surface biochemistry (von Kügelgen et al., 2024), and facilitate communication with neighbouring cells (Asmar et al., 2017). In microorganisms, cell envelopes are absolutely critical for survival, facilitating diverse functions such as attachment to surfaces (Ochner et al., 2024; O’Toole & Kolter, 1998), biofilm formation (Böhning et al., 2024), and mediating interactions with other species (Limoli et al., 2019). The composition and architecture of every cell envelope are both adapted to the specific environmental conditions that each microbe inhabits, reflecting the evolutionary history of the particular microorganism (Johnston et al., 2024).

Bacterial cell envelopes are complex, multi-layered assemblies that are typically classified into two groups based on their ability to retain Gram stain: Gram-positive bacteria, which are usually monoderm, and Gram-negative bacteria, which are usually diderm (Shugar & Baranowska, 1954), with a few notable exceptions (Baumeister & Kübler, 1978; Beaud Benyahia et al., 2024; Bharat et al., 2023; Daffé & Marrakchi, 2019). Both these major groups of bacteria have an inner membrane (IM) or cell membrane, which is a phospholipid bilayer that encloses the cellular contents. In Gram-positive bacteria, the IM is covered by a thick peptidoglycan (PG) layer, often referred to as the cell wall. In contrast, Gram-negative bacteria have a thinner PG layer and are enveloped by a second lipid bilayer called the outer membrane (OM), which is often decorated by lipopolysaccharides (LPS) (Viljoen et al., 2020). Many Gram-negative and Gram-positive bacteria have a proteinaceous, paracrystalline surface (S)-layer as the outermost coat of their cell envelope (von Kügelgen et al., 2023; Lanzoni-Mangutchi et al., 2022; Sogues et al., 2023; Bharat et al., 2017; Fagan & Fairweather, 2014; Baranova et al., 2012; Gambelli et al., 2024; Fioravanti et al., 2019; Wang et al., 2023). S-layers are composed of repeating units of S-layer proteins (SLPs) that form a two-dimensional lattice encapsulating the cell (Johnston et al., 2024; Sleytr et al., 2014). They are found in many bacteria and in almost all archaea, where they play key roles in maintaining cell shape, defence, and pathogenicity (Albers & Meyer, 2011; Bharat et al., 2021; Fagan & Fairweather, 2014).

Corynebacteria are a part of the family *Mycobacteriaceae* (based on the Genome Taxonomy Database, GTDB (Parks et al., 2022)) and typically feature an atypical cell envelope that differs both from classical Gram-positive and Gram-negative bacteria. While classified as Gram-positive due to their staining behaviour, corynebacteria are diderm, and their outermost membrane, known as the mycomembrane (MM), is markedly different from typical Gram-negative OMs (Daffé & Marrakchi, 2019; Minnikin, 1982). This distinctive cell envelope contains an IM, a cell wall containing PG and arabinogalactan (AG) layers, and the MM. The inner leaflet of the MM contains long fatty acyl chains called mycolic acids that are esterified to AG (Daffé & Marrakchi, 2019), while the outer leaflet of the MM is composed of various lipids including glycolipids, phospholipids and lipoglycans (Chiaradia et al., 2017). The covalent attachment of the mycolic acids to AG makes the MM less fluid than usual Gram-negative OMs, making their biochemistry and microbiology unique (Brown et al., 2023; Liu et al., 1995). Corynebacteria include the heavily-studied *Corynebacterium glutamicum*, which is extensively used for industrial scale amino acid production (Schallmey et al., 2014). Moreover, the list of MM-containing organisms with the same atypical cell envelope include several biomedically significant species such as *Corynebacterium diphtheriae, Mycobacterium tuberculosis*, and *Mycobacterium leprae*. In summary, although bacterial species harbouring MM-containing cell envelopes are of high importance in both biotechnology and medicine, a detailed understanding of their cell envelopes remains under-explored at the ultrastructural and molecular levels.

Previously, direct visualisation of the MM was provided by cryo-electron microscopy of vitreous sections (CEMOVIS) and electron cryomicroscopy (cryo-EM) of whole cells, which demonstrated the diderm nature of the corynebacterial cell envelope (Hoffmann et al., 2008; Sani et al., 2010; Zuber et al., 2008). Building on these foundational studies, we have used *C. glutamicum* as a model for MM-containing organisms to perform high-resolution characterisation of this unusual cell envelope. We used electron cryotomography (cryo-ET) and cryo-EM of focused ion beam-milled (FIB) cells to assign different layers of the cell envelope to cryo-EM density observed in our images. Our data shows a variable cell surface architecture of *C. glutamicum*, featuring an S-layer with a distinctive repeating pattern decorating the MM in a patchy manner. To further characterise this S-layer, we isolated and solved its structure to 3.1 Å resolution using single particle cryo-EM. Our structure shows that the S-layer is composed of repeating hexameric units of the PS2 protein that form a lattice, which partially coats the cells. By combining our S-layer structure with high-resolution cryo-ET of the cell envelope and bioinformatics analyses, we provide further clues regarding the MM-anchoring mechanisms of the S-layer and offer insights into its conservation and evolution in corynebacteria. Overall, our results offer a detailed molecular-level characterisation of the corynebacterial cell surface, bridging past work and laying a strong foundation for future research on this atypical cell surface, which is of significant biotechnological and biomedical importance.

## Results

### The cell envelope of *C. glutamicum*

To examine the cell envelope of *C. glutamicum* in detail, we performed cryo-EM of the *C. glutamicum* strain 541 (ATCC-13058). *C. glutamicum* is a rod-shaped bacterium approximately 0.7 μm in diameter and 1-2.5 μm in length (Ray et al., 2022). Its thickness presents a challenge for acquiring high-contrast images of the cell surface using traditional cryo-EM techniques (Fig. S1A). To overcome this limitation, we employed FIB milling to create thin sections of the cells, which allowed us to obtain high-resolution images with enhanced contrast of the cell envelope. Using this approach, we imaged multiple *C. glutamicum* cells in FIB-milled lamellae, which was facilitated by the tendency of *C. glutamicum* cells to cluster on cryo-EM grids (Fig. S1B). We acquired both tilt series of the lamellae to achieve a three-dimensional (3D) understanding of the cell envelope, as well as two-dimensional (2D) projection images to obtain high contrast images for morphological analysis, to specifically aid comparison with previous work (Figs. 1A-B, S1C-D and Table S2). We observed *C. glutamicum* cells at various stages of the cell cycle, including single cells and dividing cells with visible septa (Fig. 1A and S1C). Both in tomograms and in single 2D images, interior details of the cells were discernible, including ribosomes, phosphate granules, and a nucleoid region, in addition to the layers of the cell envelope (Figs. 1A-B, S1C-D and Movies S1-2).

**Figure 1.**
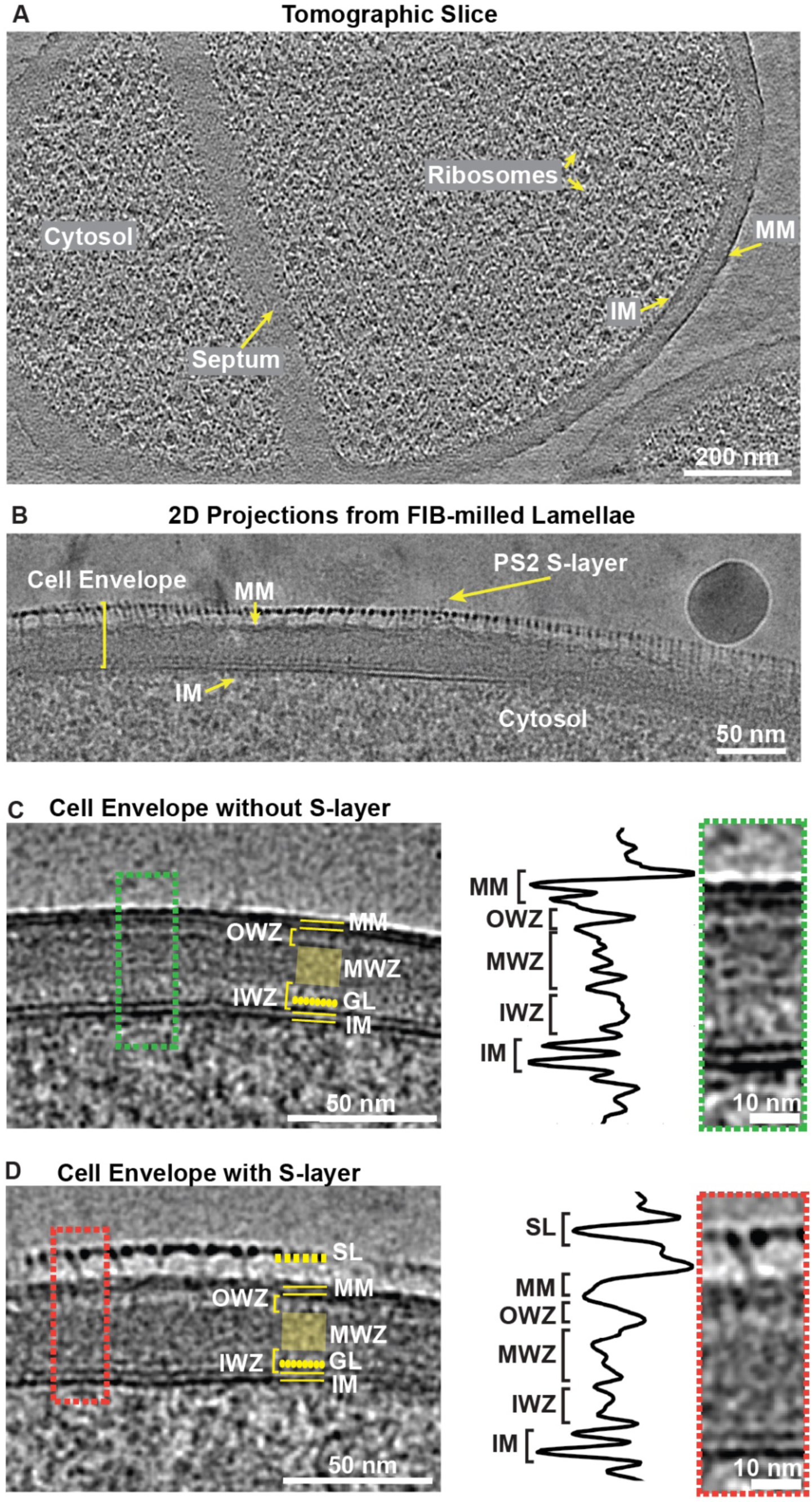
Ultrastructural investigations of the *C. glutamicum* cell envelope. **A)** A slice from a tomogram of a FIB-milled, dividing *C. glutamicum* cell. Ten slices were averaged and bandpass-filtered to boost contrast. **B)** Cryo-EM image of a FIB-milled *C. glutamicum* cell. The S-layer decorates the cell in a patchy manner (see also Fig. S1). **C)** Two-dimensional images of the FIB-milled cells without or **D)** with S-layers. The labelled features are: SL (surface layer), MM (mycomembrane), OWZ (outer wall zone), MWZ (medial wall zone), IWZ (inner wall zone), GL (granular layer), and IM (inner membrane). Line profiles were plotted from the cropped 2D projection images shown in the green (naked envelopes) and red boxes (S-layer coated envelopes). The images were Gaussian-filtered to enhance contrast.

At the outermost layer of the *C. glutamicum* cell envelope, a patchy, fragmented S-layer was observed (Fig. 1A-B and S1A, D). Previous studies have reported that the coverage of the S-layer on *C. glutamicum* cells depends on the carbon source in the medium and the strain used (Chami et al., 1995; Soual-Hoebeke et al., 1999). Consistent with this observed variability, we found that cells cultured on agar plates had higher S-layer coverage on the MM when compared to cells grown in liquid media (Chami et al., 1995). In all cases, we observed partially coated *C. glutamicum* cells, with many detached S-layers present around the cell. Beneath the S-layer on the cell, a high-contrast MM layer was observed (Figs. 1C-D). In 2D projection images of FIB-milled cells, the two leaflets of the MM were clearly resolved (Figs. 1C-D). In the case of cells without an S-layer, i.e., those containing a ‘naked’ MM, the MM presents itself as two continuous, uninterrupted, apposing leaflets, which are clearly separated from each other (Fig. 1C). This contrasts with S-layer-coated cell envelopes, where the two lines of density corresponding to the MM leaflets are interrupted and discontinuous (Fig. 1D). We hypothesised that this perturbation of the MM directly beneath the patchy S-layer is likely due to the interaction of the S-layer anchoring domain with the MM, as proposed previously (Bharat et al., 2021; Johnston et al., 2024). The thickness of the MM in both cell envelopes with and without S-layer was between 4-5 nm (Table S1), suggesting that the sizes and lengths of the mycolic acids and other components of the MM are not markedly different in these two cases.

Below the MM, there is a layer with comparatively weak density that we name the outer wall zone (OWZ), which is interrupted by densities emanating from the MM (Figs. 1C-D). This layer was not resolved in previous studies, and we propose that this comprises the AG layer, which is covalently connected both to the MM and PG (Daffé & Marrakchi, 2019). Moving inward from the OWZ, there is an electron-dense layer called the medial wall zone (MWZ), which was also observed previously (Zuber et al., 2008), and is predicted to constitute the PG. We found that this PG layer was 12-15 nm thick, with an amorphous appearance (Figs. 1C-D and Table S1). Below the MWZ, we observed another weak layer called the inner wall zone (IWZ), which has been proposed to resemble the periplasmic space of Gram-positive bacteria (Zuber et al., 2008). The IWZ is partly occupied by densities protruding from the inner membrane (IM) and this region is called the granular zone or granular layer (GL), located 4-5 nm from the edge of the IM. The presence of the GL is a property of Gram-positive bacterial PG and is thought to contain lipoproteins (Daffé & Marrakchi, 2019). Both the inner and outer leaflet of the IM are resolved in our images, with the thickness of the IM measured between 4-5 nm. Overall, we visualized all the postulated layers of the *C. glutamicum* cell envelope and found no significant difference between naked cell envelopes and S-layer-coated envelopes, apart from interruptions in the MM directly below the S-layer (Figs. 1C-D and Table S1).

### Cryo-EM analysis of the *C. glutamicum* PS2 S-layer

To understand how the S-layer is arranged on *C. glutamicum* cells, we sought to determine the atomic structure of the PS2 S-layer. To this end, we purified the S-layer from the *C. glutamicum* cell surface by adapting previous protocols that used sodium dodecyl sulphate (SDS) to detach the S-layer from bacterial cells (Johnston et al., 2024; Scheuring et al., 2002; von Kügelgen et al., 2023). Purified S-layers were deposited on cryo-EM grids and vitrified, using methods previously described for other S-layers (von Kügelgen et al., 2023, 2024), and specifically for the *C. glutamicum* S-layer (Johnston et al., 2024). Imaging the cryo-EM grids of purified S-layers confirmed a hexagonally-arranged lattice, as previously predicted (Fig. 2A and Table S3) (Johnston et al., 2024). To include some tilted and side views of the S-layer, in order to reduce resolution anisotropy and improve coverage of the Fourier space (Naydenova & Russo, 2017; Tan et al., 2017), high-throughput data collection was performed with a 30° tilt of the specimen stage (Materials and Methods). The resulting data was used to obtain a cryo-EM reconstruction at 3.1 Å global resolution (Figs. 2B and S2A-B), which allowed us to build an atomic model of the *C. glutamicum* PS2 S-layer (Figs. 2C, S2C and Table S3).

**Figure 2.**
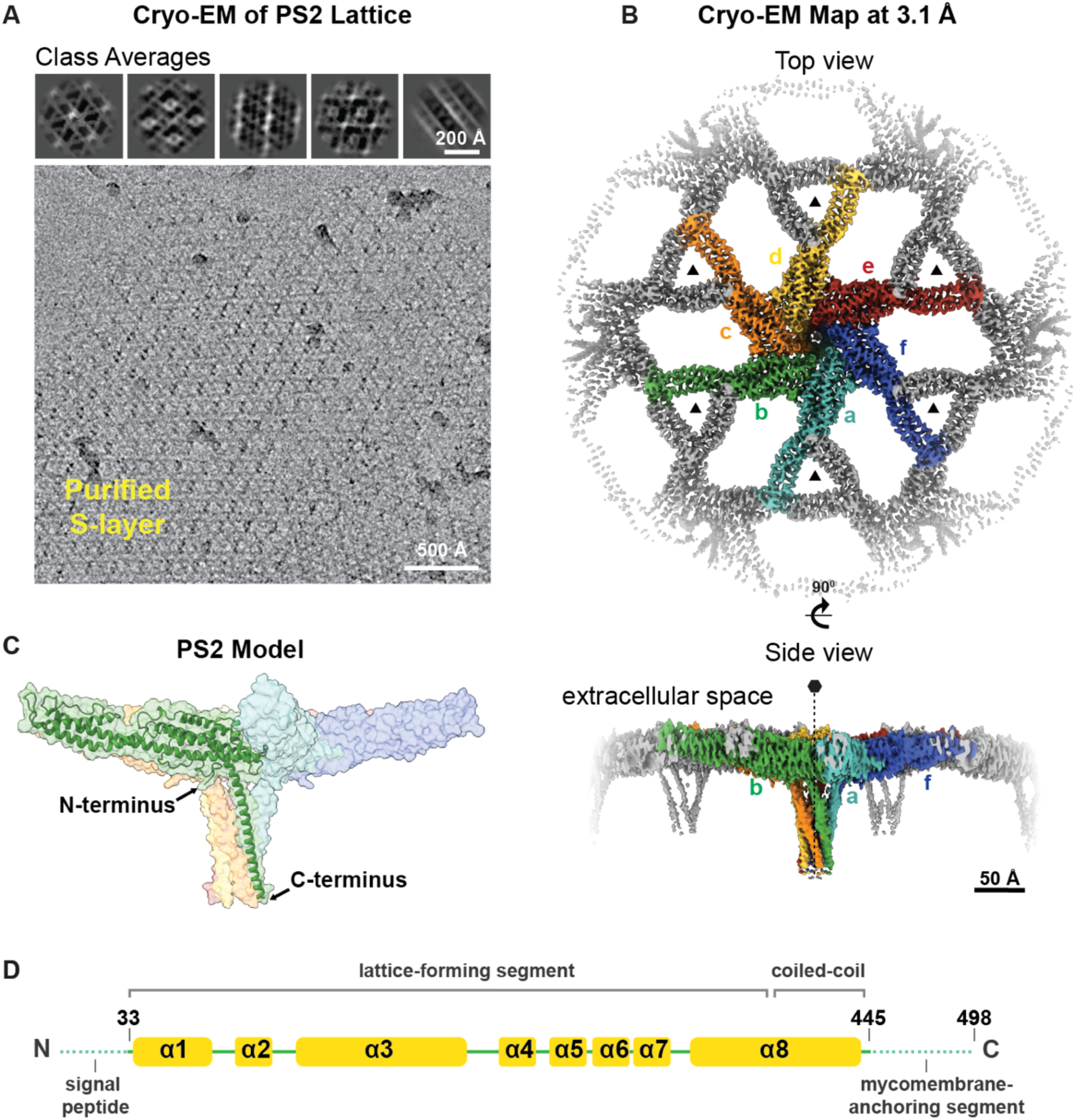
Cryo-EM structure of the PS2 S-layer of *C. glutamicum*. **A)** A cryo-EM image of the PS2 S-layer is shown, together with representative 2D classes (top). The hexagonal pattern of the PS2 S-layer is visible in the images. **B)** Cryo-EM map of the PS2 S-layer at 3.1 Å global resolution (see also Fig. S2). Both top and side views of the map are shown at a contour level 5.6 0 away from the mean value. Each PS2 monomer is coloured separately in the hexameric unit, while the rest of the volume is coloured grey. **C)** The PS2 atomic model. The N-terminal part of the protein is involved in lattice-formation, while the C-terminal part of the protein forms a coiled-coil segment that protrudes from the lattice to anchor the S-layer to the MM. **D)** A schematic cartoon of the secondary structure elements in PS2, which is composed of α-helices denoted as α1-α8. Residues 33-445 are built into the density. PS2 amino acid residues that were not built into the density are shown with a dashed line. These unmodeled residues correspond to the N-terminal region of PS2, which carries a signal peptide for its translocation, and the C-terminal regions, which contain the MM-binding residues.

Our atomic model of the S-layer lattice shows that the S-layer is formed solely by the PS2 protein, with no additional density observed for other molecules or post-translational modifications, which is consistent with previous results (Peyret et al., 1993). The PS2 protein is primarily composed of α-helices (with some loop residues), which is unusual for bacterial and archaeal SLPs structurally characterised thus far (Figs. 2D and S2C) (Bharat et al., 2017; Gambelli et al., 2022; Sogues et al., 2023; von Kügelgen et al., 2021, 2023, 2024), with the exception of the SLP of *Clostridioides difficile* (Lanzoni-Mangutchi et al., 2022). Amino acid residues 33-445 could be reliably modelled into the cryo-EM map, which were sufficient to visualise the S-layer lattice contacts (Figs. 2C-D and S2D). The missing N-terminal residues of PS2 (residues 1-32) are predicted to form a signal peptide for Sec-dependent secretion and are presumably cleaved off and not present in the mature SLP that forms the S-layer lattice. Indeed, between residues 30 and 31, there is a predicted cleavage site for signal peptidase I, which cleaves the translocated preproteins from the membrane upon their translocation (Auclair et al., 2012). At the other end of the PS2 protein sequence, the missing C-terminal residues (446-498) were previously shown to be important for the cell envelope anchoring (Chami et al., 1997) and are also predicted by bioinformatic analyses to form an anchoring segment on cells (Johnston et al., 2024). Due to detergent (SDS) treatment during S-layer purification, this anchoring segment was likely detached from the MM and remained too flexible to be resolved in our cryo-EM analysis. Instead, weak and disordered density was observed in the cryo-EM map for this part of the PS2 protein (Figs. 2D and S2E).

### The hexagonal PS2 array on the MM

The extended lattice of PS2 is formed by strong interactions between the PS2 monomers, mediated by two major symmetry-related interfaces (Fig. 3A and Table S3). The PS2 monomers assemble into tight hexamers within the lattice, which in turn are linked to each other through prominent trimeric interfaces, which are repeated throughout the lattice (Figs. 3A and S3A). The hexameric linkages are primarily established by residues 90-110 and 300-445, located around the centre of the hexamer and in the coiled-coil anchoring segment. Each hexameric unit is connected to neighbouring hexamers through trimeric interfaces, involving residues 194-204 and 213-235. Both the trimeric and hexameric interfaces are formed by a combination of hydrophilic and hydrophobic interactions, including salt-bridges and π-stacking of aromatic side chains observed in the atomic model (Figs. 3B-C).

**Figure 3.**
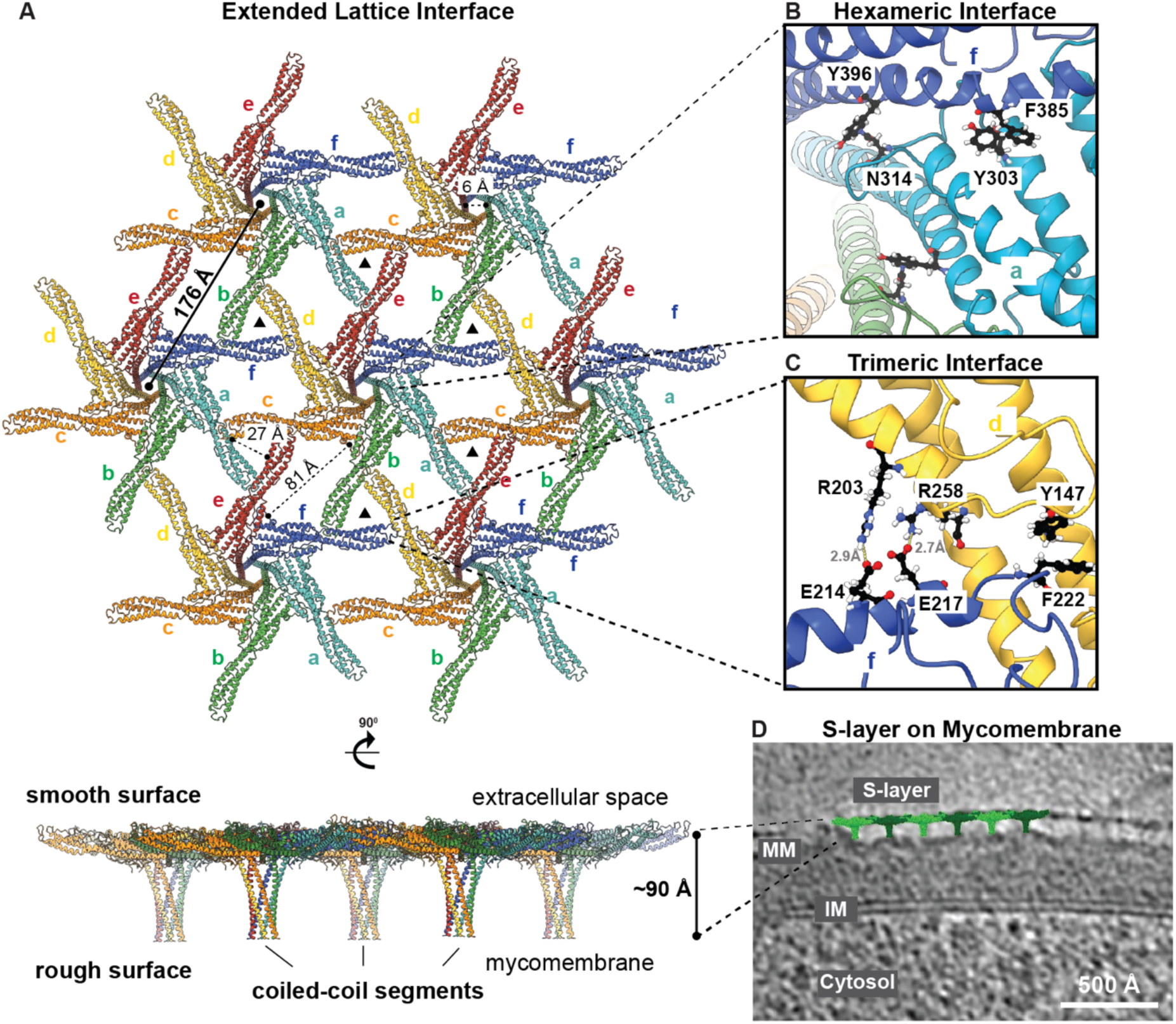
The PS2 S-layer Lattice. **A)** The atomic model of the PS2 lattice is shown in top and side views. PS2 hexamers are repeated in the 2D sheet with a lattice constant of 176 Å. Each PS2 monomer is coloured and labelled separately within the hexamer, which repeats throughout the lattice (with pore dimensions highlighted). The side view of the lattice shows the smooth surface facing the extracellular space and rough surface with coiled-coil segments protruding towards the MM. **B, C)** Interactions at the **B)** hexameric and **C)** trimeric interfaces are stabilised by both hydrophilic and hydrophobic interactions. **D)** A cell envelope tomogram (see also Fig. 1) was used to overlay the cryo-EM structure of the S-layer onto the *in situ* S-layer tomographic density. A single slice of the tomogram is shown, with the overlayed PS2 S-layer lattice in green and the tomogram in grayscale. The S-layer (Surface layer), MM (Mycomembrane), IM (Inner membrane), and Cytosol are labelled.

At the sequence level, the PS2 protein is enriched in acidic amino acid residues, giving it an overall negative charge, with an estimated isoelectric point of 4.25 (Fig. S3B-C). Consistent with this negative charge, we observed putative cationic densities at various locations along the PS2 sequence in the cryo-EM map, which are surrounded and stabilized by negatively charged amino acid residues (Figs. S3D-F). The identity of these cations cannot be ascertained at the current resolution of our cryo-EM map; however, previous studies on other bacterial S-layers suggest that they may correspond to calcium (Baranova et al., 2012; Herdman et al., 2022; Sogues et al., 2023). These cations may further stabilize the lattice, similar to other S-layers where cations were found to be essential for lattice formation (Baranova et al., 2012; Herdman et al., 2022; Sogues et al., 2023; von Kügelgen et al., 2021). Additionally, we observed one relatively larger unidentified density, surrounded by hydrophobic side-chains, which may correspond to SDS, used during the solubilization of the S-layer from the MM (Fig. S3G).

At the overall S-layer level, each PS2 protein monomer interacts with four other monomers to form the intricate lattice. Specifically, each monomer interacts with two other monomers via the hexameric interface and two others via the trimeric interface, creating a highly interconnected arrangement (Fig. 3A-C). The lattice is formed by repetition of hexameric units at a distance of 176 Å (Fig. 3A), which closely matches the lattice constant previously predicted for this S-layer (Johnston et al., 2024). The combination of hexameric and trimeric interfaces results in varying pores sizes of 6 Å, 27 Å, and 81 Å within the lattice (Fig. 3A). Some of these pores are relatively large, and are reminiscent of the porous S-layer of *Deinococcus radiodurans*, which is also patchy on the cell surface (von Kügelgen et al., 2023), suggesting that the *C. glutamicum* S-layer may not function as a molecular sieve or a barrier with a protective role.

Moving from the plane of the lattice toward the MM, the anchoring segment of PS2 consists of canonical, parallel, hexameric coiled-coils that extend inward from the S-layer lattice toward the cell. This arrangement results in a smooth extracellular side of the S-layer, while the side facing MM is rough, containing “spikes” formed by the hexameric coiled-coils (Figs. 3A, D). When viewed from the side, the purified S-layer has a thickness of 91 Å, consistent with thickness of ∼98 Å observed *in situ* in FIB-milled data from the *C. glutamicum* cell envelope (Figs. 1, 3A, 3D and Table S1). The atomic model of the S-layer lattice, when overlaid with a slice through a cell envelope tomogram of *C. glutamicum*, shows a good agreement, with a tight match to the lattice thickness and the anchoring coiled-coil densities (Fig. 3D), demonstrating that the SDS treatment did not denature the PS2 protein or disrupt the S-layer. Our overlay further shows that the tip of the coiled-coil anchoring segment, resolved in our cryo-EM map and included in the atomic model, ends just before touching the MM. This observation is consistent with the disordered C-terminal residues missing from our density (Fig. S2E and S3H), strongly suggesting that the C-terminal residues directly interact with the MM. A slight wavy pattern is observed on the MM at the bases of the PS2 coiled-coil segments (shown in Fig. 1), appearing as interruptions in the outer leaflet of the MM, further supporting the idea that the anchoring segments intimately interact with the MM.

### Conservation of the PS2 protein across corynebacterial cell surfaces

Upon characterizing the molecular structural details of the PS2 S-layer, we proceeded to investigate its presence in various species within the *Mycobacteriaceae* family. To this end, we searched for homologs of the *C. glutamicum* PS2 in the non-redundant protein sequence database (nr) at NCBI using BLAST (Camacho et al., 2009). This search yielded a total of 102 matches, almost all of which were PS2 homologs in other species of the genus *Corynebacterium*. Next, to identify highly divergent homologs of PS2 that may have remained undetected by BLAST, particularly in species outside the genus *Corynebacterium*, we conducted sensitive profile HMM-based sequence searches against representative bacterial proteomes using HHsearch (Söding, 2005; Zimmermann et al., 2018) and structural searches against the AlphaFold/UniProt50 database using Foldseek (van Kempen et al., 2024). These searches also did not find homologs of PS2 outside the *Corynebacterium* genus and further indicated that the α-helical bundle fold exhibited by PS2 is novel, strongly hinting that PS2 originated within the *Corynebacterium* genus. In fact, to date, within the MM-possessing *Mycobacteriaceae* family, S-layers have only been experimentally observed in species of the *Corynebacterium* genus, while putative SLPs, albeit of a different type containing tandem Ig-like domains, have only been bioinformatically identified in the *Nocardia* genus (Johnston et al., 2024).

Having established that the PS2 protein (and *ps2* gene) is distinct to the *Corynebacterium* genus, we next sought to scrutinise its prevalence within this group. For this, we queried a total of 2,256 proteomes of *Corynebacterium* species and their various isolates, including 73 proteomes of *C. glutamicum,* from the Refseq Genomes database and searched for PS2 using BLAST. We identified PS2 in 115 *Corynebacterium* isolates, predominantly in opportunistic pathogens isolated from human clinical and microbiome contexts, such as *C. aquatimens*, *C. aurimucosum*, *C. auris*, *C. minutissimum*, *C. singulare*, and *C. tuberculostearicum* (Supplementary information). PS2 was also found in species from environmental and biotechnological contexts, such as *C. occultum*, *C. halotolerans*, and *C. humireducens*. Interestingly, some species, such as *C. glaucum*, *C. felinum*, and *C. minutissimum*, consistently possess PS2 across all isolates, while others, including the diphtheria toxin-secreting primary pathogens *C. diphtheriae*, *C. ulcerans*, and *C. pseudotuberculosis* and their closely related species *C. rouxii*, *C. belfantii*, and *C. silvaticum* never harbour PS2. In many other species, such as *C. glutamicum*, *C. aurimucosum*, and *C. aquatimens*, the presence of PS2 varies among isolates. For instance, of the 73 *C. glutamicum* isolates in our Refseq dataset, 47 possess PS2, suggesting that the PS2 S-layer likely plays beneficial roles in *C. glutamicum* in certain conditions.

In a phylogenetic tree of *Corynebacterium* species (Fig. 4), PS2 exhibits a sparse yet widespread distribution, suggesting that the common ancestor of *Corynebacterium* likely possessed PS2, and by extension, was coated with an S-layer assembled from PS2 (Fig. 4). Notably, the phylogenetic tree also reveals a distinct separation between the PS2-lacking, diphtheria toxin-associated primary pathogens and the PS2-containing environmental and predominantly opportunistic pathogens (Fig. 4). Additionally, in a separate phylogenetic analysis we conducted on *C. glutamicum* isolates, PS2-containing and PS2-lacking isolates show a clear divergence. Many closely-related, PS2-lacking isolates cluster together in a single clade, while PS2-containing isolates are distributed across several clades. We compared the genomes of two closely related isolates—the PS2-lacking isolate CS176 and the PS2-containing isolate ATCC 31831—to identify further genetic differences related to the presence or absence of PS2. Our analysis revealed that CS176 lacks not only the gene encoding PS2 but also several adjacent genes both upstream and downstream of *ps2*. Additionally, both strains possess more than 200 distinct genes each. However, we could not establish a direct connection between these genomic differences and the presence or absence of an S-layer.

**Figure 4.**
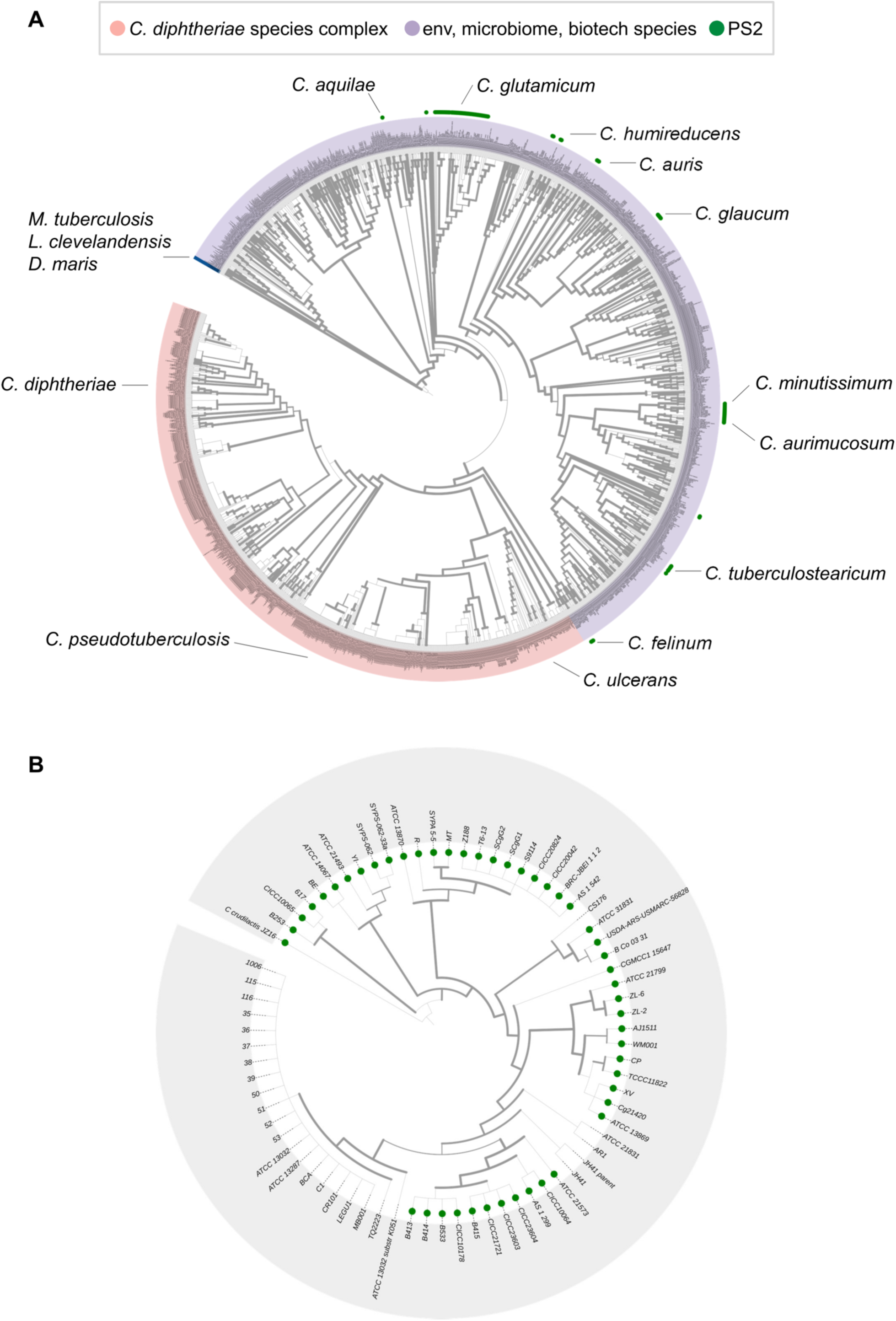
Phylogenetic Distribution of PS2. **A)** The occurrence of PS2 in species across the *Corynebacterium* genus is indicated by green dots on a GTDB species-level tree. The outer ring distinguishes diphtheria toxin-associated species (coloured light red), environmental, microbiome-associated, and opportunistic pathogenic species (light purple), and outgroup MM-containing species used to root the tree (blue; *Mycobacterium tuberculosis*, *Lawsonella clevelandensis*, and *Dietzia maris*). Some representative PS2-containing species are labelled. Line thickness represents FastTree2 Shimodaira-Hasegawa-like support values, with thicker lines indicating strong support for the branch and thinner lines suggesting that the branch is less reliable. **B)** The occurrence of PS2 in *C. glutamicum* isolates is indicated by green dots. The tree was rooted using *C. crudailactius* as the outgroup species.

At the protein sequence level, the length of PS2 from different *C. glutamicum* strains ranges from 490 to 510 amino acid residues, while the pairwise sequence identity between them ranges from 70% to 100% (Fig. S4). Variations are primarily localized to secondary structural elements involved in the formation of trimeric and hexameric interfaces in the S-layer lattice. In fact, a previous study using AFM imaging of S-layer lattices from different strains with varying PS2 sequences revealed differences in unit cell dimensions (Hansmeier et al., 2004). Beyond *C. glutamicum* strains, PS2 exhibits much greater variability, with lengths ranging from 452 to 665 residues and pairwise sequence identity ranging from 30% to 80% (Fig. S5). Despite this variability, all PS2 homologs share common sequence features, including a predicted Sec/SPI signal peptide, a coiled-coil segment, a C-terminal MM-binding hydrophobic segment, and an intrinsically disordered region between the coiled-coil and MM-binding segments. Furthermore, AlphaFold2 models of PS2 from different species reveal a high conservation of secondary structural elements, albeit with length variations, and a similar overall 3D fold (Fig. S6). This suggests that all PS2 homologs form S-layers with a similar hexagonal lattice as described in this study, but with lattice parameters that likely vary substantially.

Inspecting the variation in the PS2 amino acid sequence intimates some interesting aspects about the corynebacterial cell surface. Given that different corynebacterial species inhabit diverse environments, the varying S-layer characteristics likely play a crucial role in environmental adaptation. While the disordered region is quite variable in sequence, the length of the coiled-coil stalk and the MM-binding segment is highly conserved among PS2 homologs across species (Figs S4-S6). This is in line with the fact that the underlying cell envelope architecture, including the MM, is preserved among different *Corynebacterium* species, necessitating the conservation of the MM anchoring segments in PS2. The MM-binding segment comprises an N-terminal hydrophobic α-helix and a short C-terminal amphipathic α-helix, which includes the last residue of PS2—a phenylalanine (F) that is essentially invariant across all PS2 homologs (Figs S4-S5). The functional significance of this conserved phenylalanine residue remains unclear. However, given that the preceding residues, particularly the penultimate residue in PS2, which is typically either a proline (P) or lysine (K), are also conserved, it is tempting to speculate that these terminal residues may facilitate the sorting, export, and insertion of the protein into the MM or may play a role in ensuring the protein remains anchored in the lipid-rich MM. Notably, based on the conservation of the phenylalanine residue, we were also able to identify a few more putative MM-associated proteins in *C. glutamicum* (Fig. S7), such as an ExeM/NucH family extracellular endonuclease (NCBI: WP_003854007.1) and a prenyltransferase/squalene oxidase repeat-containing protein (WP_003854342.1).

## Discussion

In this study, we visualized the *C. glutamicum* cell envelope at high-resolution by imaging FIB-milled cells using cryo-EM and cryo-ET, allowing us to report a detailed model of the corynebacterial cell envelope (Fig. 5). Although this cell envelope had previously been visualized in impressive detail using the CEMOVIS method (Hoffmann et al., 2008; Zuber et al., 2008), our study leverages the latest cryo-ET technology, which circumvents several issues associated with CEMOVIS, most specifically sample deformation caused by the diamond knife (Al-Amoudi et al., 2005; Ochner & Bharat, 2023). We report several important characteristics of the cell envelope of *C. glutamicum* using our imaging approach (Fig. 1). We have observed a layer in the cell envelope situated just below the MM, which we have termed the OWZ. This layer forms a less dense region between the MM and MWZ layers and contains some densities that connect the MM and MWZ. We propose that this region likely corresponds to AG, which is known to be covalently linked both to PG and the MM (Daffé & Marrakchi, 2019). Below the PG layer, we observed the IWZ and GL, in line with previous observations (Figs. 1 and 5). This assignment of the densities in FIB-milled images provides an ultrastructural framework for the future study of the cell envelopes of corynebacteria and mycobacteria (i.e., members of the family *Mycobacteriaceae*), and our images will serve as a reference point for future inquiries into this topic (Fig. 5).

**Figure 5.**
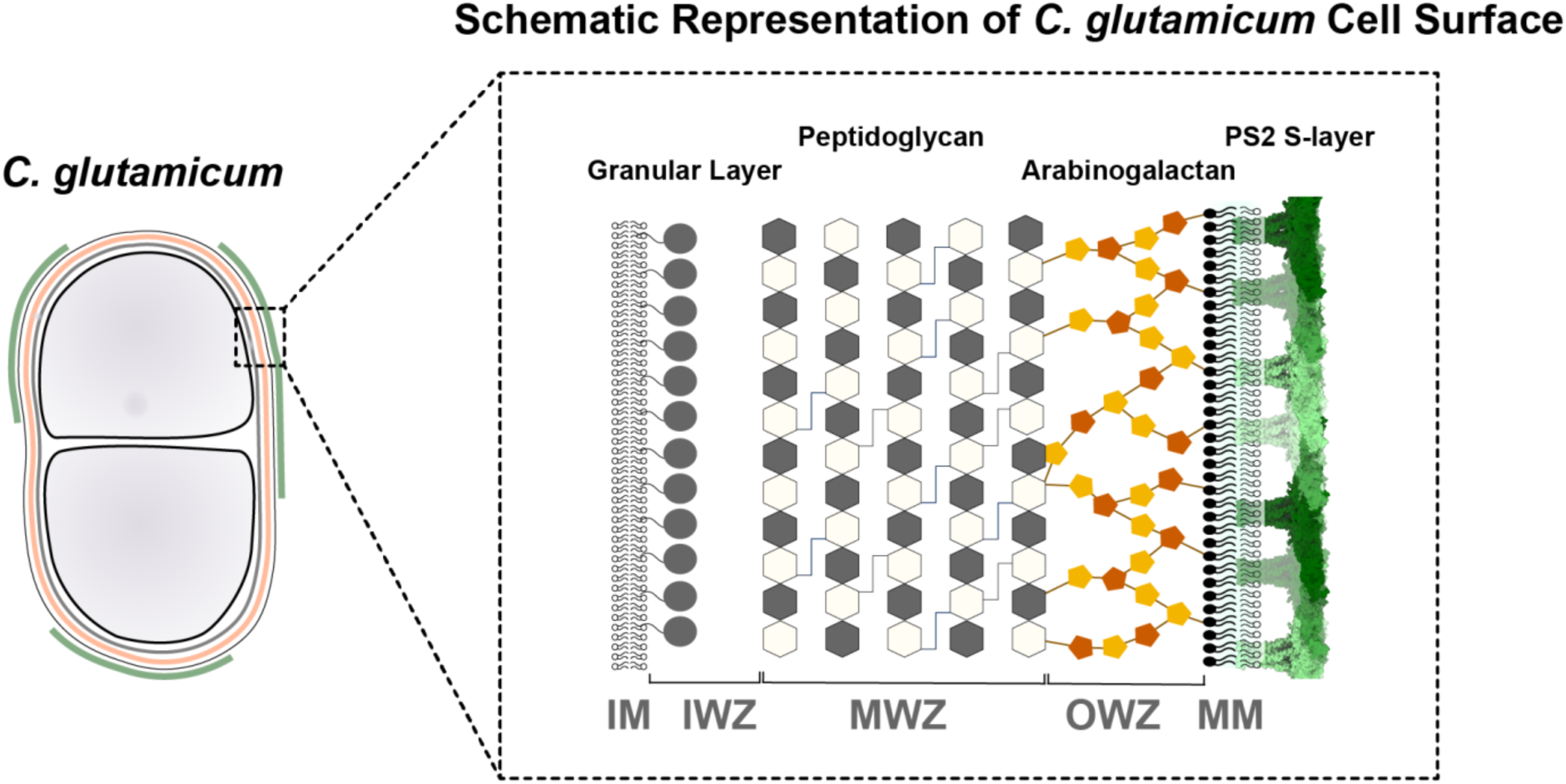
Schematic Representation of the *C. glutamicum* Cell Surface. The model depicts the layers of the *C. glutamicum* cell envelope. The S-layer forms the outermost layer, followed by the MM (mycomembrane). The MM is linked to the arabinogalactan layer (OWZ: Outer Wall Zone), which is covalently linked to peptidoglycan layer (MWZ: Medial Wall zone). Below the MWZ is the IWZ (Inner Wall Zone), which contains the granular layer. The innermost layer is the IM (inner membrane), which encloses the cellular contents.

In addition to the overall characterisation of the corynebacterial cell surface, we focused on one of the most abundant proteins on this cell surface, which also forms the outermost surface -the S-layer. We structurally characterized the PS2 S-layer of *C. glutamicum* using biochemical purification and cryo-EM structure determination (Figs. 2-3). PS2 is an SLP formed solely of α-helices, which is unusual for prokaryotic S-layers (Bharat et al., 2021; Johnston et al., 2024). PS2 possesses a C-terminal coiled-coil-containing segment that anchors it to the MM. Our analysis showed that the MM-anchoring segment, including the coiled coil, is conserved across all corynebacterial PS2 proteins, indicating its importance in anchoring the S-layer to the cell. This coiled-coil segment may function as a molecular spacer between hydrophobic MM and the hydrophilic S-layer, potentially creating an enclosed space for increasing the local concentration of certain secreted macromolecules. This arrangement could facilitate specific biochemical reactions near the cell surface, in the same manner as for other prokaryotic S-layers (von Kügelgen et al., 2021, 2024).

We have further shown that the PS2 S-layer is formed by several acidic amino acid residues (with an estimated isoelectric point of 4.25), making this an extremely hydrophilic surface modification on *C. glutamicum* cells. Given the hydrophobic nature of membranes, including the MM (Viljoen et al., 2021), the presence of the S-layer likely alters the local charge at places on the cell surface with S-layer coating, thereby significantly influencing the bacterium’s interactions with its environment. We propose that the patchy S-layer observed on *C. glutamicum* modulates the surface properties of the cell envelope in a manner similar to extended surface appendage like pili (Ochner et al., 2024) or adhesins (Melia et al., 2021). The large pores (especially the 27 Å- and 81 Å-pores) in the S-layer suggest that its role is not to protect the cells from invading molecules or phages. It is notable that in other S-layers with large pores in the lattice, such as the *D. radiodurans* S-layer made of HPI protein (von Kügelgen et al., 2023), a similarly patchy coverage of the S-layer lattice was observed, supporting this idea.

Our bioinformatic analysis of the PS2 protein sheds lights on the evolution of corynebacterial cell surfaces. Phylogenetic analysis reveals a clear distinction between the PS2-lacking, diphtheria toxin-associated primary pathogens and the PS2-containing environmental and mostly opportunistic pathogens. This suggests that primary pathogens lost the energetically costly S-layer as they adopted a pathogenic lifestyle. The conservation of the C-terminal coiled-coil and anchoring segments, compared to the relative divergence of PS2 residues involved in lattice formation, further implies that while the S-layer has diverged within corynebacteria, the MM-anchoring mechanism has likely remained the same. This suggests that the biochemistry of the MM between different organisms is largely the same, which has significant implications for our understanding of the cell surfaces of pathogens from the *Mycobacteriaceae* family, including *M. tuberculosis, M. abscessus, M. leprae*, and *C. diphteriae*. Finally and intriguingly, the conservation of specific C-terminal residues in the PS2 protein across species suggests that a precise cellular protein-targeting may be operating in *C. glutamicum* and other MM-containing species, which could be exploited for the development of therapeutic strategies against pathogenic species. This observation also holds exciting potential for future cell biology studies on these organisms.

---

**Figure S1.**
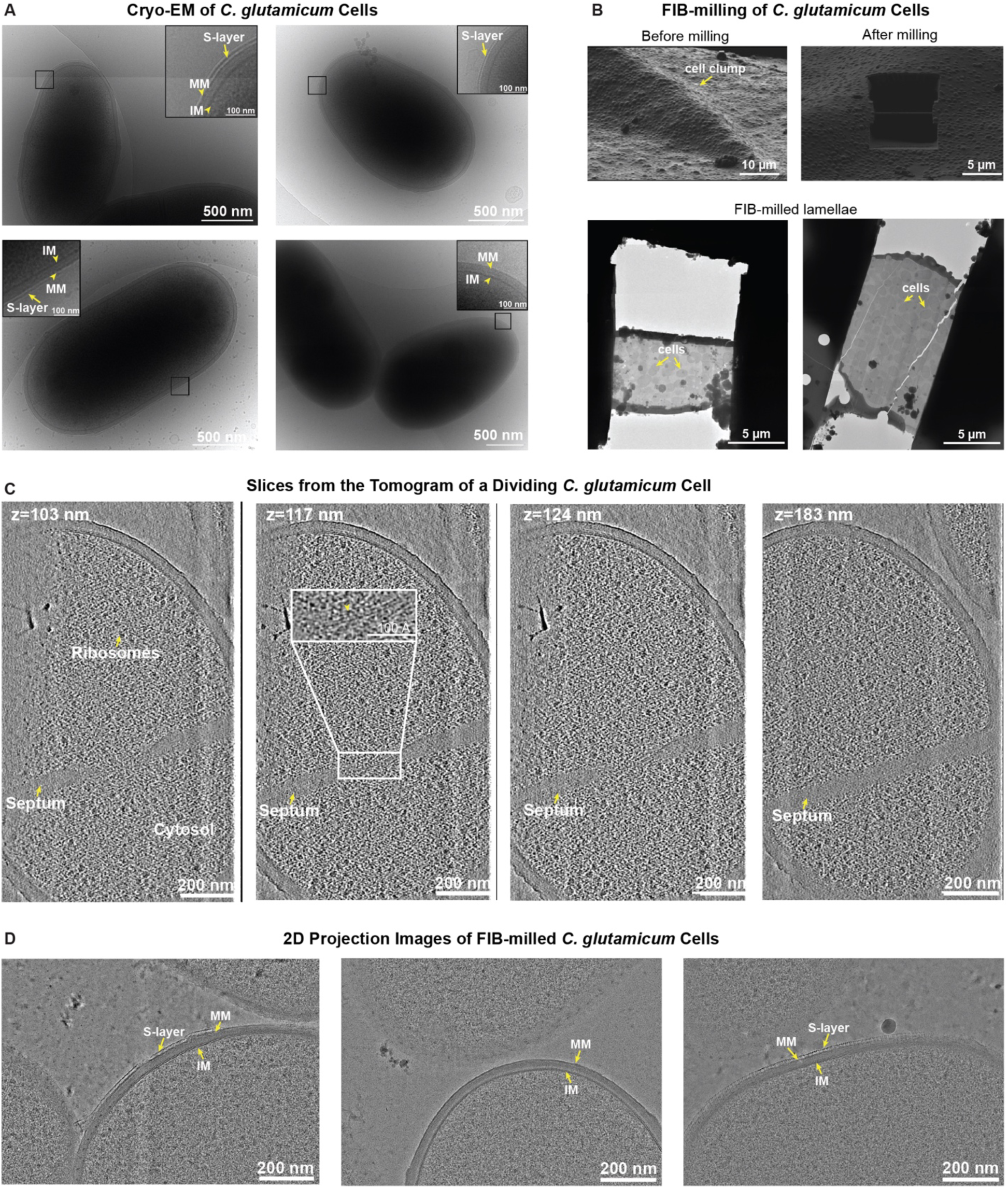
Cryo-FIB milling of *C. glutamicum* cells. **A)** Cryo-EM images of *C. glutamicum* cells deposited on grids without FIB milling. The S-layer decorates the *C. glutamicum* cell envelope in a patchy manner. S-layer (surface layer), MM (mycomembrane), and IM (inner membrane) are marked. **B)** FIB milling of *C. glutamicum* cells. Grids made for FIB-milling contained clumps of *C. glutamicum* cells, providing several suitable areas for milling. After milling, lamellae with a 150-200 nm thickness were retained for cryo-ET investigations. Each lamella contained multiple cells suitable for imaging. **C)** Slices from a tomogram of a dividing *C. glutamicum* cell. The slices were bandpass-filtered to enhance contrast. The first slice was used as a reference point to calculate Z values (marked in nm). **D)** Two-dimensional projection images of FIB-milled *C. glutamicum* cells. The 2D projection images showed high-contrast details of the cell envelope. The S-layer, MM, and IM are marked. The images were Gaussian-filtered to enhance contrast.

**Figure S2.**
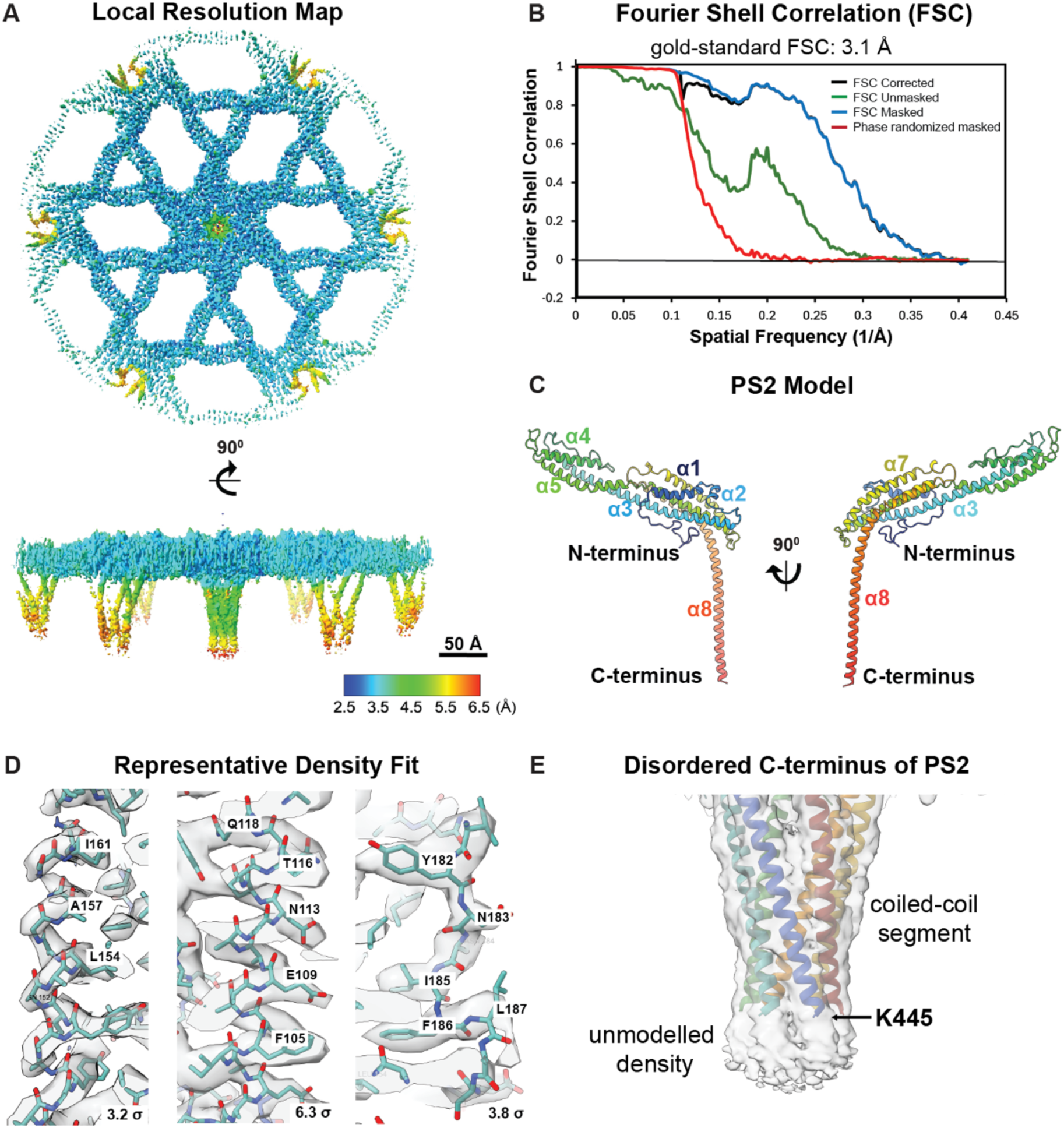
Cryo-EM analysis of the PS2 S-layer. **A)** Local resolution estimates plotted onto the cryo-EM map of the PS2 S-layer, coloured based on the local resolution values shown in the colour key (bottom right). **B)** Fourier Shell Correlation (FSC) resolution estimation of the cryo-EM map. A gold-standard FSC estimates the global resolution of the map to be 3.1 Å. **C)** PS2 atomic model shown as a ribbon diagram, coloured in a rainbow gradient with α-helices α1-α8 labelled. **D)** Representative density fits of the PS2 model. **E)** The density related to C-terminus of PS2 is disordered and could not be used for model building. The map is Gaussian-filtered with a width of 0.9 Å to illustrate the unmodeled density at the tip of coiled-coil segment.

**Figure S3.**
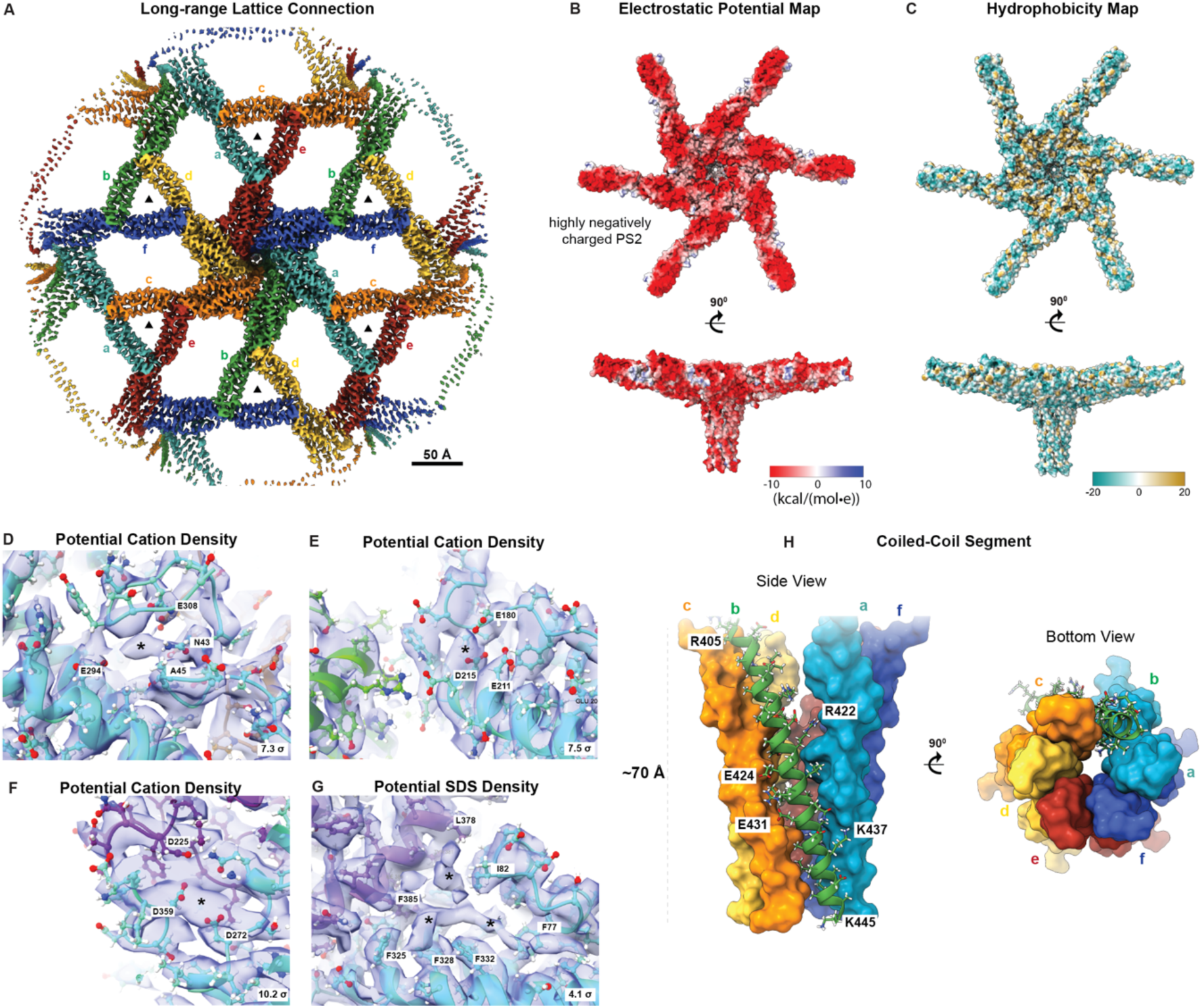
Features of the PS2 S-layer lattice. **A)** The cryo-EM map of the PS2 S-layer is shown in colour to illustrate the long-range lattice connections. Each colour represents a different monomer of PS2, labelled from ‘a’ to ‘f’. **B)** Electrostatic potential map of the PS2 hexamer. **C)** Hydrophobicity map of the PS2 hexamer. **D, E, F)** Putative densities corresponding to cations and **G)** SDS detergent molecules are shown, with the respective sigma values of the maps shown in the bottom right. The potential densities are denoted with an “*”, and the surrounding residues also labelled. **H)** The coiled-coil segment (residues 405-445) is shown in side view (left) and bottom view (right).

**Figure S4.**
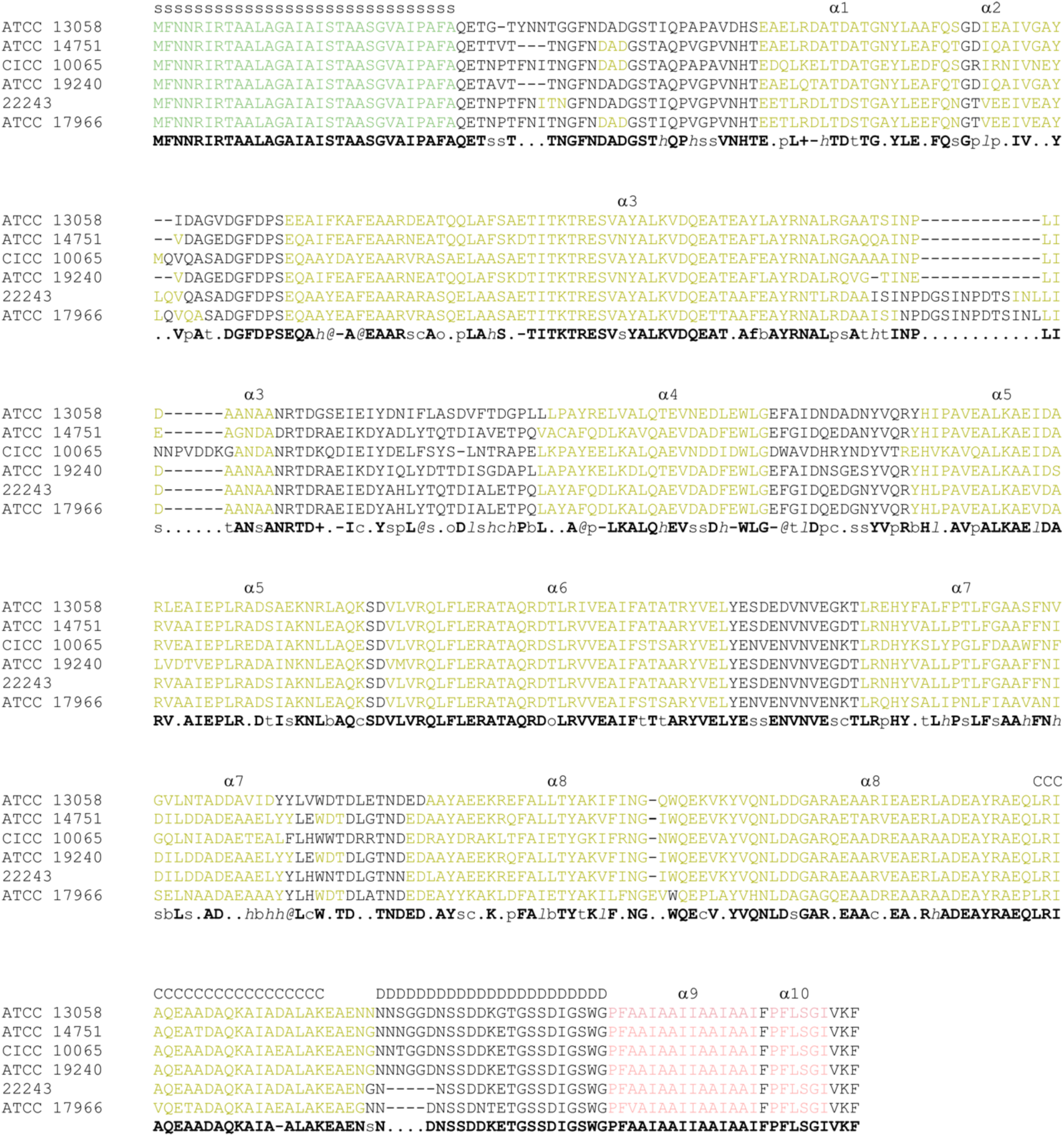
Multiple sequence alignment of PS2 from representative *C. glutamicum* isolates. The alignment of PS2 sequences from the following strains is shown: ATCC 13058 (this study), ATCC 14751 (NCBI: AAS20296.1), CICC10065 (WP_040967778.1), ATCC 19240 (AAS20313.1), 22243 (AAS20307.1), and ATCC 17966 (AAS20302.1). Signal peptides in the sequences are highlighted in green, α-helices in yellow-green, and the MM-binding segment in light red. Coiled-coil (C) and intrinsically disordered (D) regions are indicated. Secondary structure was assigned based on our experimental model of *C. glutamicum* PS2 and AlphaFold2 models. The alignment was computed using PROMALS3D. In the consensus line, conserved amino acids are represented by bold and uppercase letters, with the following annotations: aliphatic (I, V, L): l; aromatic (Y, H, W, F): @; hydrophobic (W, F, Y, M, L, I, V, A, C, T, H): h; alcohol (S, T): o; polar residues (D, E, H, K, N, Q, R, S, T): p; tiny (A, G, C, S): t; small (A, G, C, S, V, N, D, T, P): s; bulky residues (E, F, I, K, L, M, Q, R, W, Y): b; positively charged (K, R, H): +; negatively charged (D, E): -; charged (D, E, K, R, H): c.

**Figure S5.**
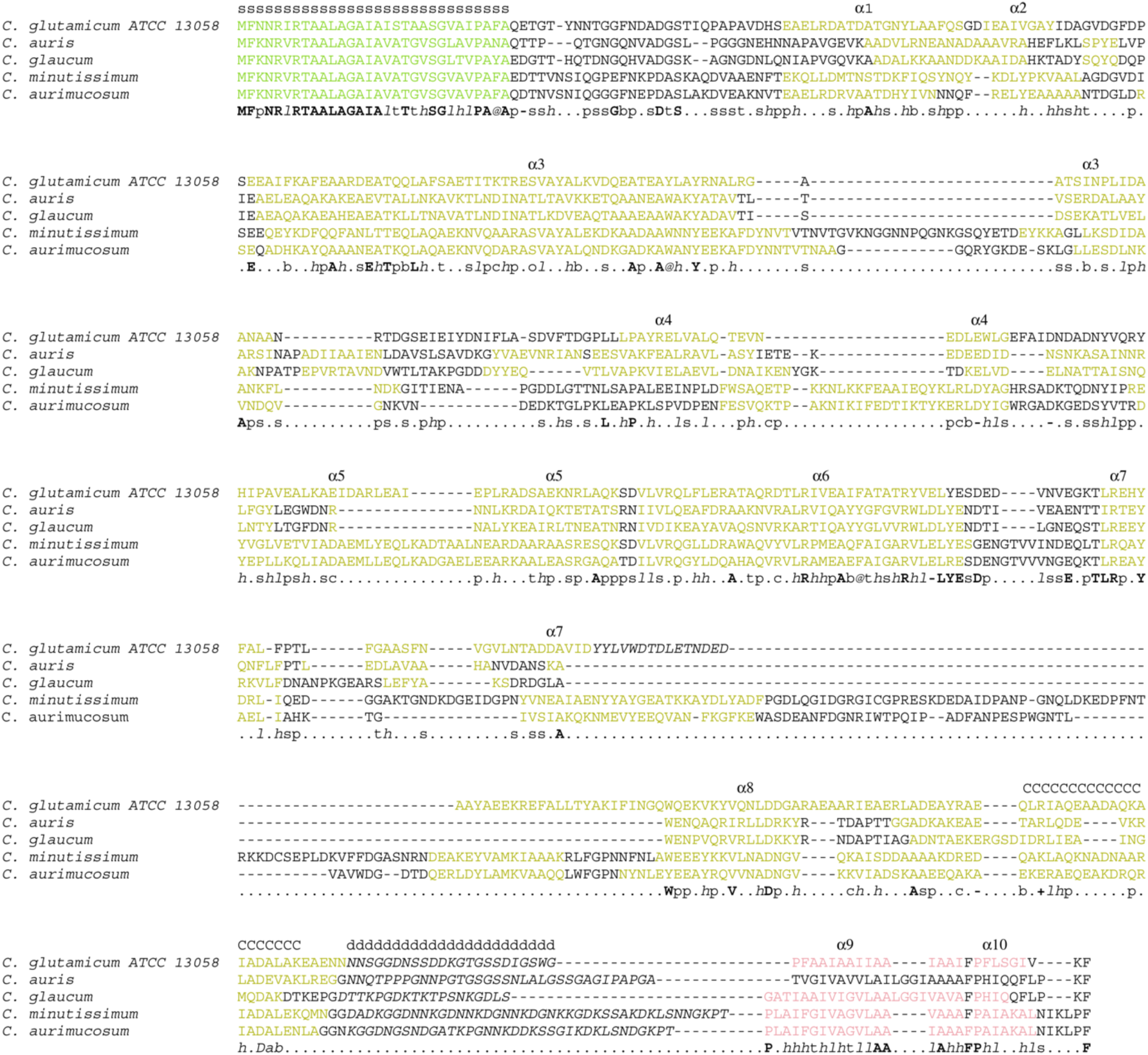
Multiple sequence alignment of PS2 from representative *Corynebacterium* species. The alignment of PS2 sequences from the following species is shown: *C. glutamicum* ATCC 13058, *C. auris*DSM 44122 (WP_290341829.1), *C. glaucum* DSM 30827 (WP_095660674.1), *C. minutissimum* NCTC10289 (WP_115020885.1), and *C. aurimucosum* UMB1300 (WP_102234280.1). The sequences are annotated as in Fig. S4.

**Figure S6.**
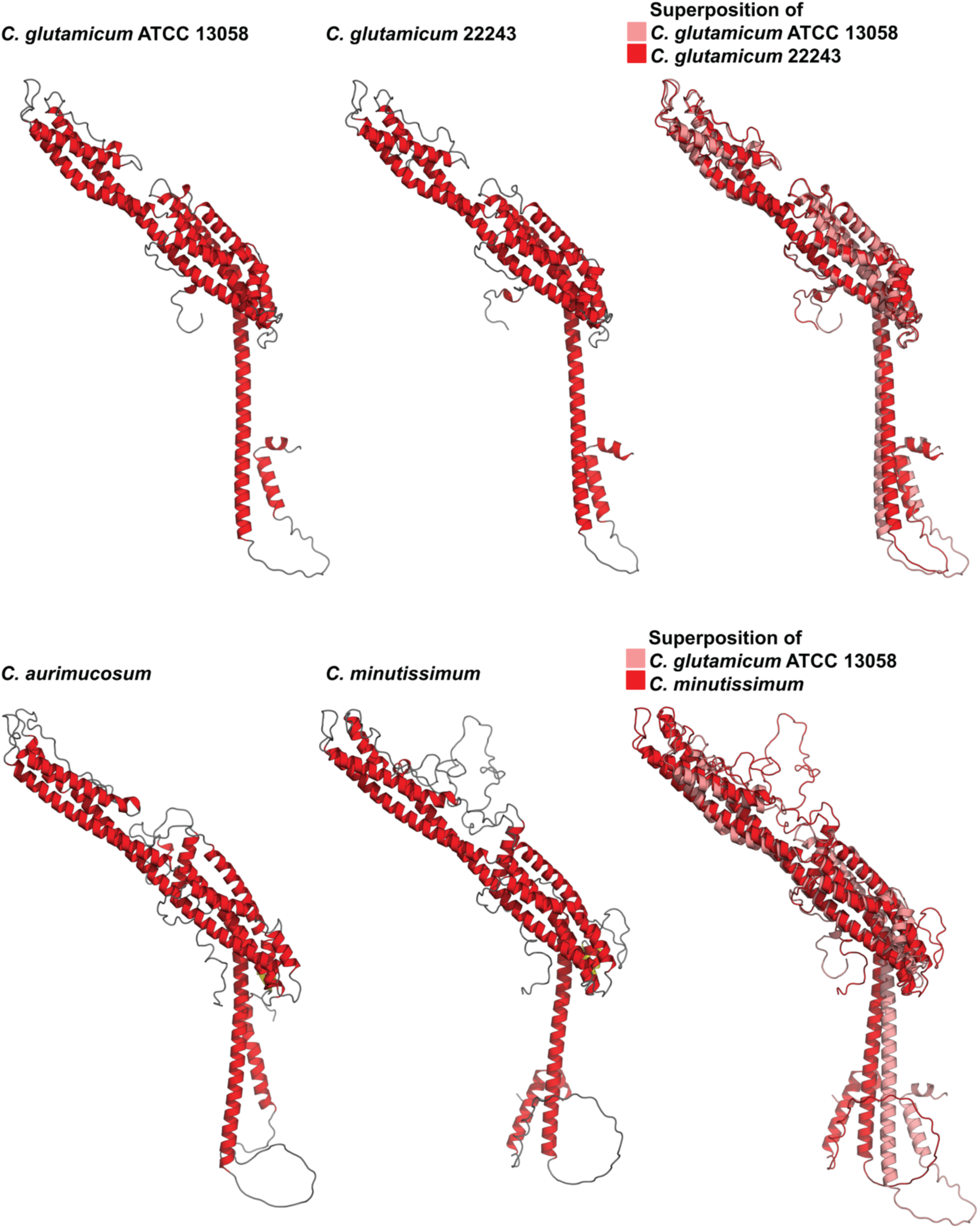
AlphaFold2-predicted monomeric structures of PS2 from representative species. In all structures, α-helices are coloured red. In the superimposed structures, one of the chains is shown in light red.

**Figure S7.**
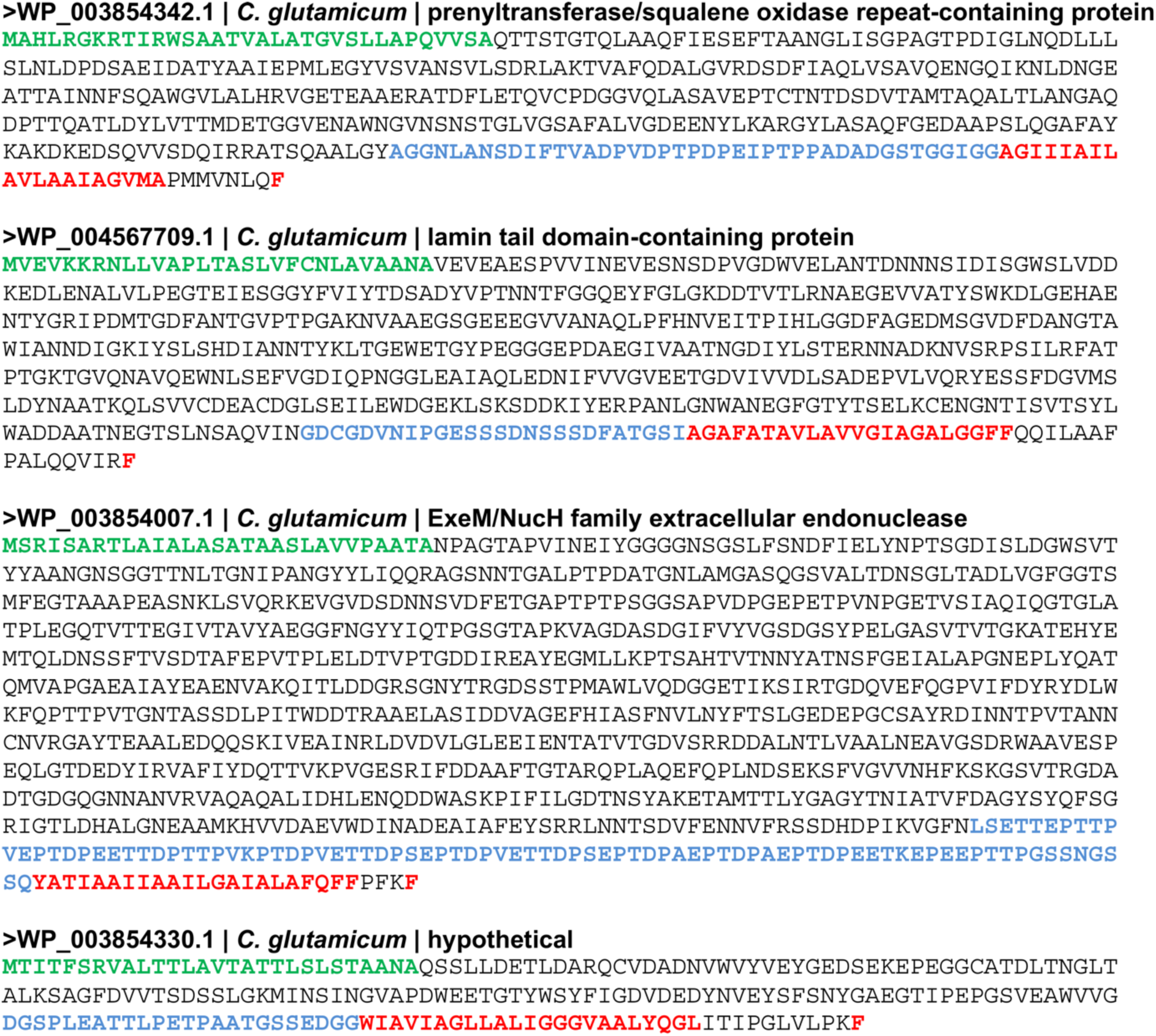
Putative MM-binding cell surface proteins in *C. glutamicum*. Predicted signal peptides are coloured green, intrinsically disordered regions are coloured blue, and the putative MM-binding segments are coloured red.

## Supplementary Tables

**Table S1.** Dimensions of cell envelope layers of *C. glutamicum*.

|  | **without S-layer** | **with S-layer** |
| --- | --- | --- |
| **S-layer** | - | 9.8 nm |
| **MM** | 5.1 nm | 4.7 nm |
| **OWZ** | 6.8 nm | 6 nm |
| **MWZ** | 15 nm | 12.7 nm |
| **IWZ** | 9.8 nm | 9.4 nm |
| **GL** | 4.7 nm | 4.3 nm |
| **IM** | 5 nm | 4.7 nm |

Measurements were performed using the plot profiles averaged over 21 nm. Apart from the membranes, measurements were performed by measuring the peak-to-peak distances in the profiles. The membrane dimensions were measured directly from the micrographs. While all measurements are consistent with previous results (Zuber et al., 2008), the IWZ is significantly thinner in our images, which could be due to strain differences.

**Table S2.**
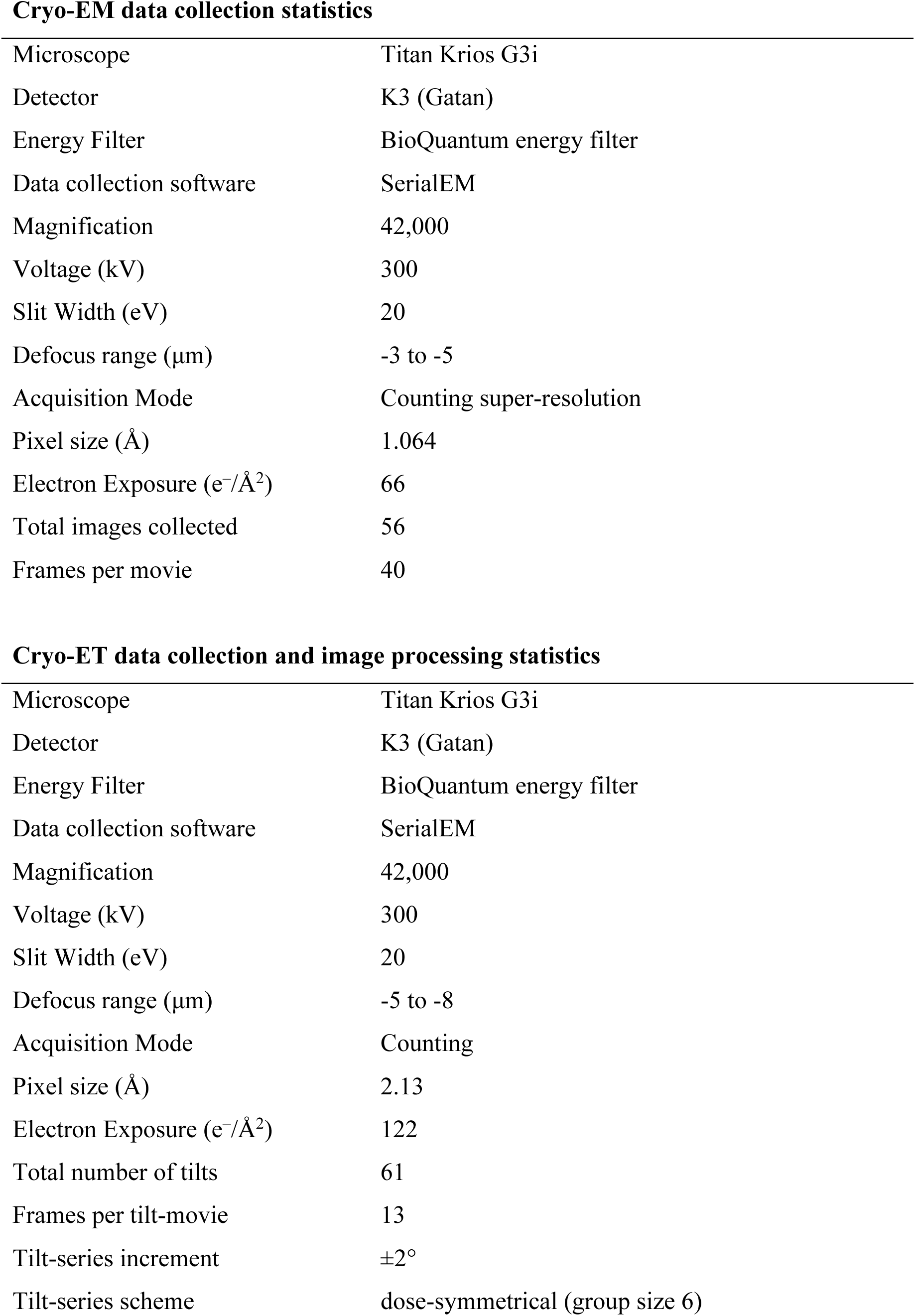
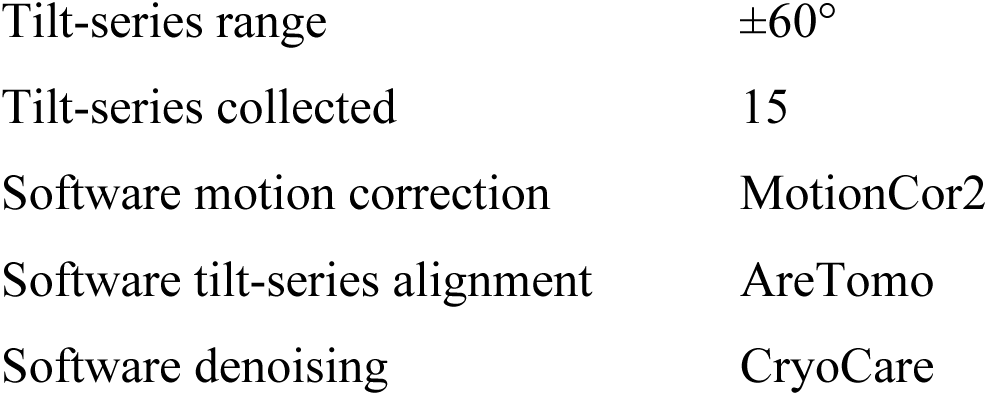
Parameters for cryo-EM and cryo-ET data collection from FIB-milled cells.

**Cryo-EM data collection statistics**
|  |  |
| --- | --- |
| Microscope | Titan Krios G3i |
| Detector | K3 (Gatan) |
| Energy Filter | BioQuantum energy filter |
| Data collection software | SerialEM |
| Magnification | 42,000 |
| Voltage (kV) | 300 |
| Slit Width (eV) | 20 |
| Defocus range ($\mu\text{m}$) | -3 to -5 |
| Acquisition Mode | Counting super-resolution |
| Pixel size ($\text{\AA}$) | 1.064 |
| Electron Exposure ($\text{e}^-/\text{\AA}^2$) | 66 |
| Total images collected | 56 |
| Frames per movie | 40 |
| **Cryo-ET data collection and image processing statistics** |  |
| Microscope | Titan Krios G3i |
| Detector | K3 (Gatan) |
| Energy Filter | BioQuantum energy filter |
| Data collection software | SerialEM |
| Magnification | 42,000 |
| Voltage (kV) | 300 |
| Slit Width (eV) | 20 |
| Defocus range ($\mu\text{m}$) | -5 to -8 |
| Acquisition Mode | Counting |
| Pixel size ($\text{\AA}$) | 2.13 |
| Electron Exposure ($\text{e}^-/\text{\AA}^2$) | 122 |
| Total number of tilts | 61 |
| Frames per tilt-movie | 13 |
| Tilt-series increment | $\pm 2^\circ$ |
| Tilt-series scheme | dose-symmetrical (group size 6) |
| Tilt-series range | $\pm 60^\circ$ |
| Tilt-series collected | 15 |
| Software motion correction | MotionCor2 |
| Software tilt-series alignment | AreTomo |
| Software denoising | CryoCare |

**Table S3.** Cryo-EM data collection, image processing and model refinement statistics of the purified PS2 S-layer.

| **Data Collection parameters of the purified PS2 S-layer** |  |
| --- | --- |
| Microscope | Titan Krios G4 |
| Detector | Falcon 4i (ThermoFisher Inc.) |
| Energy Filter | Selectrix X |
| Data collection software | EPU (ThermoFisher Inc.) |
| Magnification | 105,000 |
| Voltage (kV) | 300 |
| Slit Width (eV) | 10 |
| Defocus range ($\mu\text{m}$) | -1.6 to -2.2 |
| Acquisition Mode | Counting super-resolution |
| Pixel size ($\text{\AA}$) | 0.611 |
| Electron dose ($\text{e}^-/\text{\AA}^2$) | 50 |
| Total movies collected | 2,235 |
| Frames per movie | 40 |
| **Data Processing statistics of the purified PS2 S-layer** |  |
| Software used | CryoSPARC v4.5.3 |
| Initial particle images | 208,763 |
| Final particle images | 50,974 |
| Pre-cropped box size (pixels) | 720 x 720 x 720 |
| Final box size (pixels) | 360 x 360 x 360 |
| Pixel size final reconstruction ($\text{\AA}$) | 1.222 |
| Symmetry imposed | C6 |
| Map resolution ($\text{\AA}$) | 3.1 |
| FSC threshold | 0.143 |
| Map Sharpening B-factor ($\text{\AA}^2$) | -56 |
| **Model Refinement of PS2 S-layer** |  |
| Software | Phenix |
| Model resolution (Å) | 3.8 |
| FSC threshold | 0.5 |
| Model composition: |  |
| Residue range | 33-445 |
| Domain Range | A-R |
| Protein residues | 7434 |
| B factors (Å<sup>2</sup>) |  |
| Protein | 70.07 |
| Rms deviations: |  |
| Bond lengths (Å) | 0.003 (0) |
| Bond angles (°) | 0.560 (54) |
| MolProbity score | 1.47 |
| Clash score | 4.44 |
| Poor rotamers (%) | 0.00 |
| Cβ outliers (%) | 0.00 |
| CABLAM outliers (%) | 1.47 |
| Ramachandran plot: |  |
| Favoured (%) | 96.35 |
| Allowed (%) | 3.65 |
| Disallowed (%) | 0.00 |
| Rama-Z (Z-score, RSMD): |  |
| whole (N= 7398) | 1.06 (0.10) |
| helix (N= 4986) | 1.78 (0.07) |
| sheet (N= 0) | --- (---) |
| loop (N= 2412) | -1.67 (0.13) |

## Supplementary Movie Legends

**Movie S1. Cryo-ET of a FIB-milled *C. glutamicum* cell.**

A cryo-electron tomogram of FIB-milled *C. glutamicum* cell is shown, with the mycomembrane and inner membrane marked.

**Movie S2. Cryo-ET of a FIB-milled *C. glutamicum* cell.**

A cryo-electron tomogram of different FIB-milled *C. glutamicum* cell is shown, with the mycomembrane, inner membrane and the S-layer marked.

**Movie S3. Cryo-EM structure of the PS2 S-layer.**

The cryo-EM map and atomic structure of the *C. glutamicum* PS2 S-layer is shown. Different views of the PS2 monomer and the S-layer lattice are shown with text annotations.

## Supplementary information

**Sequence_list.csv**

A list of sequences on the PS2 protein provided in CSV format.

## Materials and Methods

### *C. glutamicum* microbiology

*Corynebacterium glutamicum* 541 (ATCC-13058) cells were cultured in liquid BE (beef extract-peptone) media at 30 °C with shaking at 180 rpm (revolutions per minute) until an OD_600_ (optical density measured at 600 nm) of 1.2 was reached. For culturing on solid media, cells were inoculated on BE agar plates (1.5 % w/v agar) and incubated at 30 °C for 18 hours.

### Purification of the PS2 S-layer

Purification of the PS2 S-layer was performed by adapting a previously used protocol (Johnston et al., 2024; von Kügelgen et al., 2023). *C. glutamicum* cells were grown until late-log phase and harvested by centrifugation at 4,000 rcf (relative centrifugal force) at 4 °C for 20 minutes. The pellet of a two litre-culture was resuspended in 100 mL lysis buffer (50 mM HEPES(4-(2-hydroxyethyl)-1-piperazineethanesulfonic acid)/NaOH pH=7.5, 150 mM NaCl, 1 mM MgCl_2_, 2 mM CaCl_2_, 50 µg/mL DNaseI, 0.2 mM TCEP (tris(2-carboxyethyl)phosphine), 2 cOmplete Protease Inhibitor tablets (Roche)). Next, SDS (sodium dodecyl sulphate) was added dropwise to the cell suspension to a final concentration of 2% (w/v). The resulting cell suspension was incubated for 70 minutes at room temperature (21 °C) and sonicated at 40% amplitude for 5 seconds in an ice bath. Cell debris was spun down at 4,000 rcf for 15 minutes at 20 °C and the supernatant was spun down once more at 35,000 rcf for 25 minutes at 20 °C. The pellet was washed with 1 mL wash buffer (50 mM HEPES/NaOH pH=7.5, 150 mM NaCl, 1 mM MgCl_2,_ 2 mM CaCl_2_) and spun down at 20,000 rcf for 15 minutes at 4 °C. The final pellet was resuspended in 120 µL of wash buffer for cryo-EM grid preparation.

### Cryo-EM grid preparation

Cryo-EM grids were prepared by adapting a previously established workflow for S-layers (Johnston et al., 2024; von Kügelgen et al., 2023). Briefly, 3.5 µL of resuspended sample was applied to a freshly glow discharged Quantifoil R2/2 Cu/Rh 200 mesh grid and plunge-frozen into liquid ethane maintained at -178 °C, using Vitrobot Mark IV after a wait time of 40 seconds, with -5 blot force and 2.5 seconds blot time. For grids containing single cells for cryoEM imaging: cells were grown in BE media until late-log phase and 3.5 µL cell suspension was applied to a freshly glow discharged Quantifoil R3.5/1 Cu/Rh 200 mesh grids and plunge-frozen using a Vitrobot Mark IV after a wait time of 40 seconds with -8 blot force and 2.5 seconds blot time. For cryo-FIB milling: cells were grown in BE media until late-log phase and spun down and resuspended in BE media supplemented with 2.5% glycerol, to an OD_600_ of 30. The resulting concentrated sample was applied (3.5 µL) to Quantifoil R1/4 SiO_2_/Au 200 mesh grids and plunge-frozen into liquid ethane using a Vitrobot Mark IV after wait time of 3 minutes with -7 blot force and 13 seconds of blot time.

### Cryo-FIB milling

Cryo-FIB milling of the *C. glutamicum* cells was performed using an Aquilos2 dual beam FIB/SEM (focused ion beam/scanning electron microscope, ThermoFisher Inc.). After loading the frozen specimen into the microscope, the grids were sputter coated with metallic platinum (0.1 mbar; 30 mA, 1V, 30 seconds), then mapped using the MAPs software (v. 3.27; ThermoFisher Inc). A coating of organic platinum was then applied via gas injection system following previous reports (Lam & Villa, 2021) for 20 seconds, followed by a final sputter-coat, using the same settings as previously. AutoTEM (v. 2.4.3; ThermoFisher Inc) was set to rough mill the lamellae in three steps with currents of 0.5 nA, 0.3 nA and 0.1 nA and with milling angle of 10°. Lamellae polishing was performed at 50 pA and then finally at 30 pA current with a 0.5° overtilt to produce lamellae with a final 100-200 nm thickness.

### Cryo-EM single-particle data collection and processing

Cryo-EM data collection was performed as described previously (Johnston et al., 2024) with a Titan Krios G4 transmission electron microscope equipped with cFEG, Falcon 4i direct electron detector and a Selectrix X energy filter with slit width 10 eV (ThermoFisher Inc.) at a nominal 105,000x magnification, resulting in a calibrated pixel size of 0.611 Å in super resolution counting mode with 30° stage tilt. The EPU software (v. 3.7.1; ThermoFisher Inc.) was used to record movies in EER format with a total electron dose of 50 e−/Å^2^ with defoci varying between -1.6 and -2.2 μm.

Cryo-EM data processing was performed using CryoSPARC v4.5.3 (Punjani et al., 2017). Movies were pre-processed using patch-motion correction and patch CTF estimation. 300 particles were manually picked and classified in 2D. With the generated 2D classes, reference-based particle picking was performed. Picked particles were inspected and filtered based on their normalized cross correlation and power threshold values. Selected particles were extracted with box size of 512 pixel^2^ and Fourier cropped to 256 pixel^2^. Particles were further classified in 2D and those particles belonging to poorly averaged classes were discarded. From the best-looking classes an ab initio volume was produced by using C6 symmetry. From the cleaned particle set, two volumes were produced with heterogenous refinement with C6 symmetry imposed. Particles belonging to the volume with higher resolution were further classified in 2D and poorly refined particles were discarded. The final particle subset was used for a homogenous refinement job which produced a cryo-EM map at 3.47 Å. This particle subset (50,974 particles) was re-extracted with 720 pixel^2^ box size (Fourier cropped to 360 pixel^2^) and local CTF refinement was performed. These steps improved the resolution of the volume to 3.35 Å with homogenous refinement. Subsequent non-uniform refinement resulted in a volume at 3.1 Å resolution. The half maps of these jobs together with the calculated FSC mask was used as input for a RELION 5.0 postprocessing, which confirmed the resolution as 3.1 Å, and the phase randomised FSC curve showed no correlation at high resolutions (Fig. S2B) (Burt et al., 2024; Chen et al., 2013).

Two-dimensional projection images of FIB-milled lamellae were collected with Titan Krios G3i transmission electron microscope (Thermo Fisher Inc.) equipped with K3 direct electron detector (Gatan) and Quantum energy filter (slit width 20 eV). Prior to data collection, the angle of lamella with respect to the stage was detected (either +10 or -10) and the specimen stage was tilted to bring the lamella perpendicular to beam path. Data collection was performed manually with the SerialEM software (Mastronarde, 2003) at a nominal 42,000x magnification, resulting in a calibrated pixel size of 1.064 Å in super resolution counting mode with 66 −/Å^2^ total electron dose and defoci varying between -3 and -5 μm. The pre-processing of the movies was performed with patch-motion correction using CryoSPARC v4.5.3 (Punjani et al., 2017). The distance measurements were performed on the 2D projection images by generating density profiles, which were calculated on rectangular selections (20 nm x 60 nm) by using FIJI (Schindelin et al., 2012).

### PS2 atomic model building and refinement

The final volume of the PS2 hexamer was sharpened in CryoSPARC with B-factor of -56 prior to model building. As an additional test, the volume was also sharpened with DeepEMhancer (Sanchez-Garcia et al., 2021) for visual inspection (not used for refinement). The exact sequence of the *ps2* gene from *C. glutamicum* 541 ATCC-13058 was obtained by whole genome sequencing of extracted genomic DNA (shown in Figs. S4-5). A partial/truncated model was automatically generated using the sharpened map by running ModelAngelo (Jamali et al., 2024). Combining the output from ModelAngelo with the AlphaFold2 prediction, a composite starting model was generated (Jumper et al., 2021). Following initial rigid body fitting, the rest of the model was inspected and manually corrected in Coot 0.9.8 (Emsley & Cowtan, 2004). The sequence register was confirmed based on side chains in the best resolved regions.

The model building was initially completed for a monomer manually using ISOLDE 1.8 and Coot followed by real space refinement performed in PHENIX 1.20.1 (Croll, 2018; Emsley & Cowtan, 2004; Liebschner et al., 2019). Next, four interacting monomers (in the hexameric and trimeric interfaces) were added to the model and refined together to ensure proper refinement of the inter-subunit interfaces. In the final rounds of real space refinement, a hexameric model was refined with non-crystallographic symmetry constraints by PHENIX real space refinement. The atomic model validation was performed within the PHENIX suite. ChimeraX was used for data visualization (Meng et al., 2023).

### Cryo-ET data collection and processing

Cryo-ET data was collected as previously described (Von Kügelgen et al., 2020). Briefly, data collection was performed with Titan Krios G3i transmission electron microscope (Thermo Fisher Inc.) equipped with a K3 direct electron detector (Gatan) and Quantum energy filter (slit width 20 eV). Prior to data collection, the angle of lamellae with respect to the stage was detected and the stage was tilted to have the lamellae perpendicular to beam path. Tilt series were then acquired with the SerialEM software (Mastronarde, 2003) using a dose-symmetric scheme (Hagen et al., 2017) at a nominal 42,000x magnification (calibrated pixel size of 2.13 Å) in counting mode with defoci between -5 to -8 µm, ±60° oscillation and 2° tilt increment with a total dose of 122 e−/Å^2^.

Movies of each tilt image were aligned with MotionCor2 (Zheng et al., 2017). Tilt series alignment and tomogram generation were performed with AreTomo (S. Zheng et al., 2022). Tomograms were denoised with CryoCare (Buchholz et al., 2019). Further visual inspection was performed within the IMOD package (Kremer et al., 1996). The overlay of the S-layer atomic model in tomograms was done using ArtiaX in ChimeraX (Ermel et al., 2022; Meng et al., 2023).

### Bioinformatic analysis

All sequence similarity searches were performed using the BLAST web server at NCBI (Camacho et al., 2009) and HHsearch in the MPI Bioinformatics Toolkit (Söding, 2005; Zimmermann et al., 2018). The searches were seeded with the PS2 protein sequence from *C. glutamicum* ATCC 13058. To assess for the prevalence of PS2 in species within the *Corynebacterium* genus, we downloaded a total of 2,315 *Corynebacterium* proteomes from the Refseq genomes database (O’Leary et al., 2016), filtering out duplicate proteomes from identical isolates to retain 2,256 unique proteomes for further analysis. These proteomes were pooled together, and a custom BLAST database was built using *makeblastdb* with default settings. We searched for the presence of the PS2 protein across *Corynebacterium* species by running BLAST with default settings, using the PS2 protein sequence from *C. glutamicum* ATCC 13058 as the query against our custom *Corynebacterium* database. Signal peptides were predicted using SignalP 6.0 (Teufel et al., 2022), and multiple sequence alignments were computed using PROMALS3D (Pei & Grishin, 2014). The phylogenetic trees of *Corynebacterium* species and *C. glutamicum* isolates, shown in Fig. 4, were generated using the de novo workflow (*de_novo_wf*) of the GTDB-Tk 2.4.0 software toolkit (Chaumeil et al., 2022) with default settings. For the tree generation, uncultured and undetermined species were omitted. The tree of *Corynebacterium* species contains 1,585 species, including three outgroup species used for rooting the tree (*Dietzia maris* IMV 195T; NCBI GCF_014144855.1, *Lawsonella clevelandensis* X1036; GCF_001293125.1, and *Mycobacterium tuberculosis* H37Rv; GCF_000195955.2). The tree of *C. glutamicum* isolates contains 73 isolates and one outgroup species used for rooting (*C. crudilactis* JZ16; GCF_001643015.1). The trees were visualized in iTOL (Letunic & Bork, 2024). Structural models of PS2 from representative species were predicted using AlphaFold v2.3.2 (Jumper et al., 2021).

## Acknowledgments

This work was supported by the Medical Research Council, as part of United Kingdom Research and Innovation (also known as UK Research and Innovation) [Programme MC_UP_1201/31 to T.A.M.B.]. T.A.M.B. and B.I. would like to thank the Human Frontier Science Program (Grant RGY0074/2021), the European Molecular Biology Organization, the Wellcome Trust (Grant 225317/Z/22/Z), the Leverhulme Trust, and the Lister Institute for Preventative Medicine for support. V.A. would like to thank Andrei Lupas for continued support and the Human Frontier Science Program (Grant RGY0074/2021). The authors would like to thank Abul Tarafder, Ido Caspy, and Jan Böhning critically reading this manuscript. We acknowledge the MRC LMB electron microscopy facility for help with sample preparation and data collection and the MRC LMB mass spectrometry facility for assistance with mass spectrometry analysis of the sample.

## Author Contributions

B.I. and T.A.M.B. designed research. B.I., A.Y., V.A., and T.A.M.B. performed research. B.I., V.A., and T.A.M.B. analysed data. B.I., V.A., and T.A.M.B. wrote the manuscript with support from A.Y.

## Data Availability

The cryo-EM density maps and atomic models for the *C. glutamicum* S-layer structure presented will be deposited at the EMDB and PDB on acceptance and released prior to publication. Please direct all requests for materials to the corresponding author T.A.M.B.

## References

Al-Amoudi, A., Studer, D., & Dubochet, J. (2005). Cutting artefacts and cutting process in vitreous sections for cryo-electron microscopy. Journal of Structural Biology, 150(1), 109–121. 10.1016/j.jsb.2005.01.003

Albers, S.-V., & Meyer, B. H. (2011). The archaeal cell envelope. Nature Reviews Microbiology, 9(6), 414–426. 10.1038/nrmicro2576

Asmar, A. T., Ferreira, J. L., Cohen, E. J., Cho, S.-H., Beeby, M., Hughes, K. T., & Collet, J.-F. (2017). Communication across the bacterial cell envelope depends on the size of the periplasm. PLOS Biology, 15(12), e2004303. 10.1371/journal.pbio.2004303

Auclair, S. M., Bhanu, M. K., & Kendall, D. A. (2012). Signal peptidase I: Cleaving the way to mature proteins. Protein Science : A Publication of the Protein Society, 21(1), 13–25. 10.1002/pro.757

Baranova, E., Fronzes, R., Garcia-Pino, A., Van Gerven, N., Papapostolou, D., Péhau-Arnaudet, G., Pardon, E., Steyaert, J., Howorka, S., & Remaut, H. (2012). SbsB structure and lattice reconstruction unveil Ca2+ triggered S-layer assembly. Nature, 487(7405), 119–122. 10.1038/nature11155

Baumeister, W., & Kübler, O. (1978). Topographic study of the cell surface of micrococcus radiodurans. Proceedings of the National Academy of Sciences of the United States of America, 75(11), 5525–5528.

Beaud Benyahia, B., Taib, N., Beloin, C., & Gribaldo, S. (2024). Terrabacteria: Redefining bacterial envelope diversity, biogenesis and evolution. Nature Reviews Microbiology, 1–16. 10.1038/s41579-024-01088-0

Bharat, T. A. M., Kureisaite-Ciziene, D., Hardy, G. G., Yu, E. W., Devant, J. M., Hagen, W. J. H., Brun, Y. V., Briggs, J. A. G., & Löwe, J. (2017). Structure of the hexagonal surface layer on Caulobacter crescentus cells. Nature Microbiology, 2, 17059. 10.1038/nmicrobiol.2017.59

Bharat, T. A. M., Tocheva, E. I., & Alva, V. (2023). The cell envelope architecture of Deinococcus: HPI forms the S-layer and SlpA tethers the outer membrane to peptidoglycan. Proceedings of the National Academy of Sciences, 120(51), e2305338120. 10.1073/pnas.2305338120

Bharat, T. A. M., von Kügelgen, A., & Alva, V. (2021). Molecular Logic of Prokaryotic Surface Layer Structures. Trends in Microbiology, 29(5), 405–415. 10.1016/j.tim.2020.09.009

Böhning, J., Tarafder, A. K., & Bharat, T. A. M. (2024). The role of filamentous matrix molecules in shaping the architecture and emergent properties of bacterial biofilms. The Biochemical Journal, 481(4), 245–263. 10.1042/BCJ20210301

Brown, T., Chavent, M., & Im, W. (2023). Molecular Modeling and Simulation of the Mycobacterial Cell Envelope: From Individual Components to Cell Envelope Assemblies. The Journal of Physical Chemistry B, 127(51), 10941–10949. 10.1021/acs.jpcb.3c06136

Buchholz, T.-O., Krull, A., Shahidi, R., Pigino, G., Jékely, G., & Jug, F. (2019). Content-aware image restoration for electron microscopy. Methods in Cell Biology, 152, 277–289. 10.1016/bs.mcb.2019.05.001

Burt, A., Toader, B., Warshamanage, R., Von Kügelgen, A., Pyle, E., Zivanov, J., Kimanius, D., Bharat, T. A. M., & Scheres, S. H. W. (2024). An image processing pipeline for electron cryo-tomography in RELION-5. 10.1101/2024.04.26.591129

Camacho, C., Coulouris, G., Avagyan, V., Ma, N., Papadopoulos, J., Bealer, K., & Madden, T. L. (2009). BLAST+: Architecture and applications. BMC Bioinformatics, 10, 421. 10.1186/1471-2105-10-421

Cell surface patterning and morphogenesis: Biogenesis of a periodic surface array during Caulobacter development. (1982). The Journal of Cell Biology, 95(1), 41– 49.

Chami, M., Bayan, N., Dedieu, J.-C., Leblon, G., Shechter, E., & Gulik-Krzywicki, T. (1995). Organization of the outer layers of the cell envelope of Corynebacterium glutamicum: A combined freeze-etch electron microscopy and biochemical study. Biology of the Cell, 83(2–3), 219–229. 10.1016/0248-4900(96)81311-6

Chami, M., Bayan, N., Peyret, J. L., Gulik-Krzywicki, T., Leblon, G., & Shechter, E. (1997). The S-layer protein of Corynebacterium glutamicum is anchored to the cell wall by its C-terminal hydrophobic domain. Molecular Microbiology, 23(3), 483–492. 10.1046/j.1365-2958.1997.d01-1868.x

Chaumeil, P.-A., Mussig, A. J., Hugenholtz, P., & Parks, D. H. (2022). GTDB-Tk v2: Memory friendly classification with the genome taxonomy database. Bioinformatics (Oxford, England), 38(23), 5315–5316. 10.1093/bioinformatics/btac672

Chen, S., McMullan, G., Faruqi, A. R., Murshudov, G. N., Short, J. M., Scheres, S. H. W., & Henderson, R. (2013). High-resolution noise substitution to measure overfitting and validate resolution in 3D structure determination by single particle electron cryomicroscopy. Ultramicroscopy, 135, 24–35. 10.1016/j.ultramic.2013.06.004

Chiaradia, L., Lefebvre, C., Parra, J., Marcoux, J., Burlet-Schiltz, O., Etienne, G., Tropis, M., & Daffé, M. (2017). Dissecting the mycobacterial cell envelope and defining the composition of the native mycomembrane. Scientific Reports, 7(1), 12807. 10.1038/s41598-017-12718-4

Croll, T. I. (2018). ISOLDE: A physically realistic environment for model building into low-resolution electron-density maps. *Acta Crystallographica. Section D*, Structural Biology, 74(Pt 6), 519–530. 10.1107/S2059798318002425

Daffé, M., & Marrakchi, H. (2019). Unraveling the Structure of the Mycobacterial Envelope. Microbiology Spectrum, 7(4), 10.1128/microbiolspec.gpp3-0027-2018. https://doi.org/10.1128/microbiolspec.gpp3-0027-2018

Emsley, P., & Cowtan, K. (2004). Coot: Model-building tools for molecular graphics. Acta Crystallographica Section D: Biological Crystallography, 60(12), 2126–2132. 10.1107/S0907444904019158

Ermel, U. H., Arghittu, S. M., & Frangakis, A. S. (2022). ArtiaX: An electron tomography toolbox for the interactive handling of sub-tomograms in UCSF ChimeraX. Protein Science: A Publication of the Protein Society, 31(12), e4472. 10.1002/pro.4472

Fagan, R. P., & Fairweather, N. F. (2014). Biogenesis and functions of bacterial S-layers. Nature Reviews Microbiology, 12(3), 211–222. 10.1038/nrmicro3213

Fioravanti, A., Van Hauwermeiren, F., Van der Verren, S. E., Jonckheere, W., Goncalves, A., Pardon, E., Steyaert, J., De Greve, H., Lamkanfi, M., & Remaut, H. (2019). Structure of S-layer protein Sap reveals a mechanism for therapeutic intervention in anthrax. Nature Microbiology, 4(11), 1805–1814. 10.1038/s41564-019-0499-1

Gambelli, L., McLaren, M., Conners, R., Sanders, K., Gaines, M. C., Clark, L., Gold, V. A., Kattnig, D., Sikora, M., Hanus, C., Isupov, M. N., & Daum, B. (n.d.). Structure of the two-component S-layer of the archaeon Sulfolobus acidocaldarius. eLife, 13, e84617. 10.7554/eLife.84617

Hagen, W. J. H., Wan, W., & Briggs, J. A. G. (2017). Implementation of a cryo-electron tomography tilt-scheme optimized for high resolution subtomogram averaging. Journal of Structural Biology, 197(2), 191–198. 10.1016/j.jsb.2016.06.007

Hansmeier, N., Bartels, F. W., Ros, R., Anselmetti, D., Tauch, A., Pühler, A., & Kalinowski, J. (2004). Classification of hyper-variable Corynebacterium glutamicum surface-layer proteins by sequence analyses and atomic force microscopy. Journal of Biotechnology, 112(1–2), 177–193. 10.1016/j.jbiotec.2004.03.020

Herdman, M., von Kügelgen, A., Kureisaite-Ciziene, D., Duman, R., El Omari, K., Garman, E. F., Kjaer, A., Kolokouris, D., Löwe, J., Wagner, A., Stansfeld, P. J., & Bharat, T. A. M. (2022). High-resolution mapping of metal ions reveals principles of surface layer assembly in Caulobacter crescentus cells. Structure(London, England:1993), 30(2), 215-228.e5. 10.1016/j.str.2021.10.012

Hoffmann, C., Leis, A., Niederweis, M., Plitzko, J. M., & Engelhardt, H. (2008). Disclosure of the mycobacterial outer membrane: Cryo-electron tomography and vitreous sections reveal the lipid bilayer structure. Proceedings of the National Academy of Sciences, 105(10), 3963–3967. 10.1073/pnas.0709530105

Jamali, K., Käll, L., Zhang, R., Brown, A., Kimanius, D., & Scheres, S. H. W. (2024). Automated model building and protein identification in cryo-EM maps. Nature, 628(8007), 450–457. 10.1038/s41586-024-07215-4

Johnston, E., Isbilir, B., Alva, V., Bharat, T. A. M., & Doye, J. P. K. (2024). Punctuated and continuous structural diversity of S-layers across the prokaryotic tree of life (p. 2024.05.28.596244). bioRxiv. 10.1101/2024.05.28.596244

Jumper, J., Evans, R., Pritzel, A., Green, T., Figurnov, M., Ronneberger, O., Tunyasuvunakool, K., Bates, R., Žídek, A., Potapenko, A., Bridgland, A., Meyer, C., Kohl, S. A. A., Ballard, A. J., Cowie, A., Romera-Paredes, B., Nikolov, S., Jain, R., Adler, J., … Hassabis, D. (2021). Highly accurate protein structure prediction with AlphaFold. Nature, 596(7873), 583–589. 10.1038/s41586-021-03819-2

Kremer, J. R., Mastronarde, D. N., & McIntosh, J. R. (1996). Computer Visualization of Three-Dimensional Image Data Using IMOD. Journal of Structural Biology, 116(1), 71–76. 10.1006/jsbi.1996.0013

Lanzoni-Mangutchi, P., Banerji, O., Wilson, J., Barwinska-Sendra, A., Kirk, J. A., Vaz, F., O’Beirne, S., Baslé, A., El Omari, K., Wagner, A., Fairweather, N. F., Douce, G. R., Bullough, P. A., Fagan, R. P., & Salgado, P. S. (2022). Structure and assembly of the S-layer in C. difficile. Nature Communications, 13(1), 970. 10.1038/s41467-022-28196-w

Letunic, I., & Bork, P. (2024). Interactive Tree of Life (iTOL) v6: Recent updates to the phylogenetic tree display and annotation tool. Nucleic Acids Research, 52(W1), W78–W82. 10.1093/nar/gkae268

Liebschner, D., Afonine, P. V., Baker, M. L., Bunkóczi, G., Chen, V. B., Croll, T. I., Hintze, B., Hung, L.-W., Jain, S., McCoy, A. J., Moriarty, N. W., Oeffner, R. D., Poon, B. K., Prisant, M. G., Read, R. J., Richardson, J. S., Richardson, D. C., Sammito, M. D., Sobolev, O. V., … Adams, P. D. (2019). Macromolecular structure determination using X-rays, neutrons and electrons: Recent developments in Phenix. Acta Crystallographica Section D: Structural Biology, 75(10), 861–877. 10.1107/S2059798319011471

Limoli, D. H., Warren, E. A., Yarrington, K. D., Donegan, N. P., Cheung, A. L., & O’Toole, G. A. (2019, November 12). Interspecies interactions induce exploratory motility in Pseudomonas aeruginosa. eLife; eLife Sciences Publications Limited. 10.7554/eLife.47365

Liu, J., Rosenberg, E. Y., & Nikaido, H. (1995). Fluidity of the lipid domain of cell wall from Mycobacterium chelonae. Proceedings of the National Academy of Sciences, 92(24), 11254–11258. 10.1073/pnas.92.24.11254

Mastronarde, D. N. (2003). SerialEM: A Program for Automated Tilt Series Acquisition on Tecnai Microscopes Using Prediction of Specimen Position. Microscopy and Microanalysis, 9(S02), 1182–1183. 10.1017/S1431927603445911

Melia, C. E., Bolla, J. R., Katharios-Lanwermeyer, S., Mihaylov, D. B., Hoffmann, P. C., Huo, J., Wozny, M. R., Elfari, L. M., Böhning, J., Morgan, A. N., Hitchman, C. J., Owens, R. J., Robinson, C. V., O’Toole, G. A., & Bharat, T. A. M. (2021). Architecture of cell–cell junctions in situ reveals a mechanism for bacterial biofilm inhibition. Proceedings of the National Academy of Sciences of the United States of America, 118(31), e2109940118. 10.1073/pnas.2109940118

Meng, E. C., Goddard, T. D., Pettersen, E. F., Couch, G. S., Pearson, Z. J., Morris, J. H., & Ferrin, T. E. (2023). UCSF ChimeraX: Tools for structure building and analysis. Protein Science : A Publication of the Protein Society, 32(11), e4792. 10.1002/pro.4792

Minnikin. (1982). Lipids: Complex lipids, their chemistry, biosynthesis and roles. In Ratledge C, Stanford JL (eds) The biology of the mycobacteria. (pp. 95–184). Academic Press, London.

Naydenova, K., & Russo, C. J. (2017). Measuring the effects of particle orientation to improve the efficiency of electron cryomicroscopy. Nature Communications, 8(1), 629. 10.1038/s41467-017-00782-3

Ochner, H., & Bharat, T. A. M. (2023). Charting the molecular landscape of the cell. Structure, 31(11), 1297–1305. 10.1016/j.str.2023.08.015

Ochner, H., Böhning, J., Wang, Z., Tarafder, A. K., Caspy, I., & Bharat, T. A. M. (2024). Structure of the Pseudomonas aeruginosa PAO1 Type IV pilus (p. 2024.04.09.588664). bioRxiv. 10.1101/2024.04.09.588664

O’Leary, N. A., Wright, M. W., Brister, J. R., Ciufo, S., Haddad, D., McVeigh, R., Rajput, B., Robbertse, B., Smith-White, B., Ako-Adjei, D., Astashyn, A., Badretdin, A., Bao, Y., Blinkova, O., Brover, V., Chetvernin, V., Choi, J., Cox, E., Ermolaeva, O., … Pruitt, K. D. (2016). Reference sequence (RefSeq) database at NCBI: Current status, taxonomic expansion, and functional annotation. Nucleic Acids Research, 44(D1), D733–745. 10.1093/nar/gkv1189

O’Toole, G. A., & Kolter, R. (1998). Flagellar and twitching motility are necessary for Pseudomonas aeruginosa biofilm development. Molecular Microbiology, 30(2), 295–304. 10.1046/j.1365-2958.1998.01062.x

Parks, D., M, C., C, R., Aj, M., Pa, C., & P, H. (2022). GTDB: An ongoing census of bacterial and archaeal diversity through a phylogenetically consistent, rank normalized and complete genome-based taxonomy. Nucleic Acids Research, 50(D1). 10.1093/nar/gkab776

Pei, J., & Grishin, N. V. (2014). PROMALS3D: Multiple protein sequence alignment enhanced with evolutionary and three-dimensional structural information. *Methods in Molecular Biology (Clifton*, N.J.), 1079, 263–271. 10.1007/978-1-62703-646-7_17

Peyret, J. L., Bayan, N., Joliff, G., Gulik-Krzywicki, T., Mathieu, L., Shechter, E., & Leblon, G. (1993). Characterization of the cspB gene encoding PS2, an ordered surface-layer protein in Corynebacterium glutamicum. Molecular Microbiology, 9(1), 97–109. 10.1111/j.1365-2958.1993.tb01672.x

Punjani, A., Rubinstein, J. L., Fleet, D. J., & Brubaker, M. A. (2017). cryoSPARC: Algorithms for rapid unsupervised cryo-EM structure determination. Nature Methods, 14(3), Article 3. 10.1038/nmeth.4169

Sanchez-Garcia, R., Gomez-Blanco, J., Cuervo, A., Carazo, J. M., Sorzano, C. O. S., & Vargas, J. (2021). DeepEMhancer: A deep learning solution for cryo-EM volume post-processing. Communications Biology, 4(1), 1–8. 10.1038/s42003-021-02399-1

Sani, M., Houben, E. N. G., Geurtsen, J., Pierson, J., De Punder, K., Van Zon, M., Wever, B., Piersma, S. R., Jiménez, C. R., Daffé, M., Appelmelk, B. J., Bitter, W., Van Der Wel, N., & Peters, P. J. (2010). Direct Visualization by Cryo-EM of the Mycobacterial Capsular Layer: A Labile Structure Containing ESX-1-Secreted Proteins. PLoS Pathogens, 6(3), e1000794. 10.1371/journal.ppat.1000794

Schallmey, M., Frunzke, J., Eggeling, L., & Marienhagen, J. (2014). Looking for the pick of the bunch: High-throughput screening of producing microorganisms with biosensors. Current Opinion in Biotechnology, 26, 148–154. 10.1016/j.copbio.2014.01.005

Scheuring, S., Stahlberg, H., Chami, M., Houssin, C., Rigaud, J.-L., & Engel, A. (2002). Charting and unzipping the surface layer of Corynebacterium glutamicum with the atomic force microscope. Molecular Microbiology, 44(3), 675–684. 10.1046/j.1365-2958.2002.02864.x

Schindelin, J., Arganda-Carreras, I., Frise, E., Kaynig, V., Longair, M., Pietzsch, T., Preibisch, S., Rueden, C., Saalfeld, S., Schmid, B., Tinevez, J.-Y., White, D. J., Hartenstein, V., Eliceiri, K., Tomancak, P., & Cardona, A. (2012). Fiji: An open-source platform for biological-image analysis. Nature Methods, 9(7), 676–682. 10.1038/nmeth.2019

Shugar, D., & Baranowska, J. (1954). Studies on the gram stain; the importance of proteins in the Gram reaction. Acta Microbiologica Polonica (1952), 3(1), 11–20.

Sleytr, U. B., Schuster, B., Egelseer, E., & Pum, D. (2014). S-layers: Principles and applications. FEMS Microbiology Reviews, 38(5), 823–864. 10.1111/1574-6976.12063

Söding, J. (2005). Protein homology detection by HMM–HMM comparison. Bioinformatics, 21(7), 951–960. 10.1093/bioinformatics/bti125

Sogues, A., Fioravanti, A., Jonckheere, W., Pardon, E., Steyaert, J., & Remaut, H. (2023). Structure and function of the EA1 surface layer of Bacillus anthracis. Nature Communications, 14(1), 7051. 10.1038/s41467-023-42826-x

Soual-Hoebeke, E., Sousa- D’Auria, C. de, Chami, M., Baucher, M.-F., Guyonvarch, A., Bayan, N., Salim, K., & Leblon, G. (1999). S-layer protein production by Corynebacterium strains is dependent on the carbon source. Microbiology, 145(12), 3399–3408. 10.1099/00221287-145-12-3399

Tan, Y. Z., Baldwin, P. R., Davis, J. H., Williamson, J. R., Potter, C. S., Carragher, B., & Lyumkis, D. (2017). Addressing preferred specimen orientation in single-particle cryo-EM through tilting. Nature Methods, 14(8), 793–796. 10.1038/nmeth.4347

Teufel, F., Almagro Armenteros, J. J., Johansen, A. R., Gíslason, M. H., Pihl, S. I., Tsirigos, K. D., Winther, O., Brunak, S., von Heijne, G., & Nielsen, H. (2022). SignalP 6.0 predicts all five types of signal peptides using protein language models. Nature Biotechnology, 40(7), 1023–1025. 10.1038/s41587-021-01156-3

van Kempen, M., Kim, S. S., Tumescheit, C., Mirdita, M., Lee, J., Gilchrist, C. L. M., Söding, J., & Steinegger, M. (2024). Fast and accurate protein structure search with Foldseek. Nature Biotechnology, 42(2), 243–246. 10.1038/s41587-023-01773-0

Viljoen, A., Foster, S. J., Fantner, G. E., Hobbs, J. K., & Dufrêne, Y. F. (2020). Scratching the Surface: Bacterial Cell Envelopes at the Nanoscale. mBio, 11(1), 10.1128/mbio.03020-19. https://doi.org/10.1128/mbio.03020-19

Viljoen, A., Räth, E., Mckinney, J. D., Fantner, G. E., & Dufrêne, Y. F. (2021). Seeing and Touching the Mycomembrane at the Nanoscale. Journal of Bacteriology, 203(10), 10.1128/jb.00547-20. https://doi.org/10.1128/jb.00547-20

von Kügelgen, A., Alva, V., & Bharat, T. A. M. (2021). Complete atomic structure of a native archaeal cell surface. Cell Reports, 37(8), 110052. 10.1016/j.celrep.2021.110052

von Kügelgen, A., Cassidy, C. K., van Dorst, S., Pagani, L. L., Batters, C., Ford, Z., Löwe, J., Alva, V., Stansfeld, P. J., & Bharat, T. A. M. (2024). Membraneless channels sieve cations in ammonia-oxidizing marine archaea. Nature, 630(8015), 230–236. 10.1038/s41586-024-07462-5

Von Kügelgen, A., Tang, H., Hardy, G. G., Kureisaite-Ciziene, D., Brun, Y. V., Stansfeld, P. J., Robinson, C. V., & Bharat, T. A. M. (2020). In Situ Structure of an Intact Lipopolysaccharide-Bound Bacterial Surface Layer. Cell, 180(2), 348–358.e15. 10.1016/j.cell.2019.12.006

von Kügelgen, A., van Dorst, S., Yamashita, K., Sexton, D. L., Tocheva, E. I., Murshudov, G., Alva, V., & Bharat, T. A. M. (2023). Interdigitated immunoglobulin arrays form the hyperstable surface layer of the extremophilic bacterium Deinococcus radiodurans. Proceedings of the National Academy of Sciences, 120(16), e2215808120. 10.1073/pnas.2215808120

Wang, H., Zhang, J., Toso, D., Liao, S., Sedighian, F., Gunsalus, R., & Zhou, Z. H. (2023). Hierarchical organization and assembly of the archaeal cell sheath from an amyloid-like protein. Nature Communications, 14(1), 6720. 10.1038/s41467-023-42368-2

Zheng, S. Q., Palovcak, E., Armache, J.-P., Verba, K. A., Cheng, Y., & Agard, D. A. (2017). MotionCor2: Anisotropic correction of beam-induced motion for improved cryo-electron microscopy. Nature Methods, 14(4), 331–332. 10.1038/nmeth.4193

Zheng, S., Wolff, G., Greenan, G., Chen, Z., Faas, F. G. A., Bárcena, M., Koster, A. J., Cheng, Y., & Agard, D. A. (2022). AreTomo: An integrated software package for automated marker-free, motion-corrected cryo-electron tomographic alignment and reconstruction. Journal of Structural Biology: X, 6, 100068. 10.1016/j.yjsbx.2022.100068

Zimmermann, L., Stephens, A., Nam, S.-Z., Rau, D., Kübler, J., Lozajic, M., Gabler, F., Söding, J., Lupas, A. N., & Alva, V. (2018). A Completely Reimplemented MPI Bioinformatics Toolkit with a New HHpred Server at its Core. Journal of Molecular Biology, 430(15), 2237–2243. 10.1016/j.jmb.2017.12.007

Zuber, B., Chami, M., Houssin, C., Dubochet, J., Griffiths, G., & Daffé, M. (2008). Direct Visualization of the Outer Membrane of Mycobacteria and Corynebacteria in Their Native State. Journal of Bacteriology, 190(16), 5672–5680. 10.1128/jb.01919-07

